# Telomeric transposons are pervasive in linear bacterial genomes

**DOI:** 10.1101/2024.11.13.623437

**Authors:** Shan-Chi Hsieh, Máté Fülöp, Richard Schargel, Michael T. Petassi, Orsolya Barabas, Joseph E. Peters

## Abstract

Eukaryotes have linear DNA and their telomeres are hotspots for transposons, which in some cases took over telomere maintenance. While many bacteria also have linear chromosomes and plasmids, no transposons were known to target bacterial telomeres. Here we show several families of independently evolved telomeric transposons in cyanobacteria and *Streptomyces*. While these elements have one specific transposon end sequence with the second boundary being the telomere, we can show they move using two transposon ends. Telomeres are transiently bridged by the telomere maintenance systems, providing a duplex substrate for mobilization of the element and the associated telomere. We identify multiple instances where telomeric transposons have replaced native telomeres, making the host cell dependent on the new telomere system for genome maintenance. This work indicates how telomeric transposons can promote gene transfer both between and within genomes, significantly influencing the evolutionary dynamics of linear genomes.

## Introduction

Linear chromosomes were once regarded as a defining feature that differentiated eukaryotes from bacteria, whose genetic material consists primarily of circular double-stranded DNA. In recent decades, many bacterial lineages were found to have linear chromosomes and plasmids, including the plant pathogen *Agrobacterium*, pathogenic spirochaetes *Borrelia*, *Streptomyces*, and some photosynthetic cyanobacteria. In all organisms, linear DNA ends pose unique challenges, as these structures require protection against various threats, including double-strand DNA break repair pathways and the end replication problem. Two mechanisms of telomere maintenance have evolved in bacteria to overcome these challenges, which are fundamentally different from the mechanism used in eukaryotes. Eukaryotic telomeres consist of repetitive sequences located at the ends of eukaryotic chromosomes, playing a crucial role in maintaining chromosomal integrity and stability. The enzyme telomerase, a specialized reverse transcriptase, is responsible for elongating telomeres and preventing erosion of these protective structures. The two types of bacterial telomere maintenance mechanisms have distinct origins^1,2^. The first type involves the action of protelomerase enzymes, which catalyze the formation of hairpin structures at the DNA ends. The second type involves the use of terminal proteins, which serve to cap the DNA ends and act as primers for DNA replication to finish replication of the lagging-strand template at linear DNA ends.

In Eukaryotes, telomeres are known to be the target of many retrotransposons which move via an RNA intermediate. Most telomeric transposons belong to the non-LTR (long terminal repeat) retrotransposons and *Penelope*-like retroelements (PLE)^3^. In some cases, telomeric transposons have evolved to become the mechanism of telomere maintenance, as found in *Drosophila*^4^.

In the current work, we reveal that specific lineages of bacteria possessing linear chromosomes also harbor telomeric transposons. These unique genetic elements found at the ends of linear chromosomes and plasmids are DNA transposons and often contain genes responsible for maintaining bacterial telomeres. We identified three distinct groups of bacterial telomeric transposons in cyanobacteria and *Streptomyces* derived from two groups of specialized DNA transposons, Tn7-like transposons, and simpler mobile elements we call TnsBC transposons. Tn7-like transposons are known for their tight control of transposition, allowing them to selectively target transposition into specific genomic sites^5^. These transposons typically contain a set of core genes, a heteromeric transposase (TnsA and TnsB), an AAA+ ATPase regulator (TnsC), and one or more target selector proteins (e.g. TniQ). TniQ is the key factor that confers site-specificity to Tn7-like transposons. TniQ proteins have evolved distinct strategies for recognizing specific target sites, utilizing diverse DNA-binding modules that are either naturally fused to the C-terminal end of TniQ or recruited by interaction with TniQ^5^. Recently, Tn7-like CRISPR-associated transposons (CASTs) which recruit nuclease-dead CRISPR-Cas systems have gained significant attention due to their potential for RNA-guided genome editing^6,7^. Several independently evolved CASTs have been characterized, referred to by the CRISPR-Cas subtype that was coopted by the transposon^8^. Transposons we describe as TnsBC elements contain a TnsB transposase and a TnsC AAA+ ATPase regulator but lack TnsA and the TniQ target site selector protein. The TnsBC elements are DNA transposons which use a replicative transposition process that is broadly similar to the process used by bacteriophage Mu to make copies of the bacteriophage genome via proteins analogous to TnsB (Mu protein MuA) and TnsC (Mu protein MuB)^9^.

Canonical DNA transposons from the Tn7 and TnsBC groups require two ends for transposition. However, we find that all three families of bacterial telomeric transposons have only one specific transposon end with the other flanking end being the telomere. We show that examples of one-ended transposons with each telomere type move similar to common DNA transposons using two transposon ends. They can move when their telomeres are bridged, such that they mobilize the telomere during transposition, making the transposons mobile telomeres. Bioinformatics and experimental data indicate multiple adaptations these transposons use to accommodate the one-end lifestyle and provide evidence that they mediate telomere replacement. Interestingly, the telomeric transposons found in cyanobacteria are from Tn7-like elements closely related to CAST elements that coopted a type V-K CRISPR-Cas system, while one group of the TnsBC telomeric transposons include type I-E CRISPR-associated transposons (I-E CAST), the first non-Tn7-like CAST^7^. This provides compelling evidence for the remarkable evolutionary plasticity of Tn7/TnsBC transposons and underscores the convergent evolution of these mobile genetic elements to a telomeric lifestyle. These findings shed new light on the complex interplay between mobile genetic elements and their host organisms and have significant implications for our understanding of bacterial genome evolution and diversity.

## Results

A previous bioinformatic analysis of Tn7-like transposons in cyanobacteria revealed three major clades: those featuring a TnsAB fusion, those with TnsA and TnsB as separate polypeptides, and those without TnsA^10^. While most of the Tn7-like transposons in cyanobacteria that lack TnsA are type V-K CAST^10^, we identified two groups that appeared to lack the gene encoding the Cas12k effector protein. One group appears to be a standard, albeit minimal, transposon, like simple insertion sequences, encoding only TnsB, TnsC, and a gene of unknown function (CP000117.1 in Fig. 1a). The second group of Tn7-like transposons that lack TnsA and Cas12k-encoding genes, conspicuously seemed to always be found near the ends of assembled chromosomes and plasmids (“PAST” in Fig. 1a). Over half of the examples in this group (15/27) have a protelomerase gene downstream of the TnsBCQ encoding operon, and in none of the examples could we identify the second end of the transposons. In addition, some transposon containing contigs end with inverted repeats downstream of the TnsBCQ operon. Based on these observations, we hypothesized that the transposons have only one end and reside near the hairpin ends of the linear cyanobacterial chromosome, which are maintained by the protelomerase they carry. This hypothesis would account for why the transposition system was encoded near the end of the contig; the inverted repeat at the end of the contig could result from a sequencing artifact as the sequencing polymerase read through the hairpin end.

**Fig. 1.**
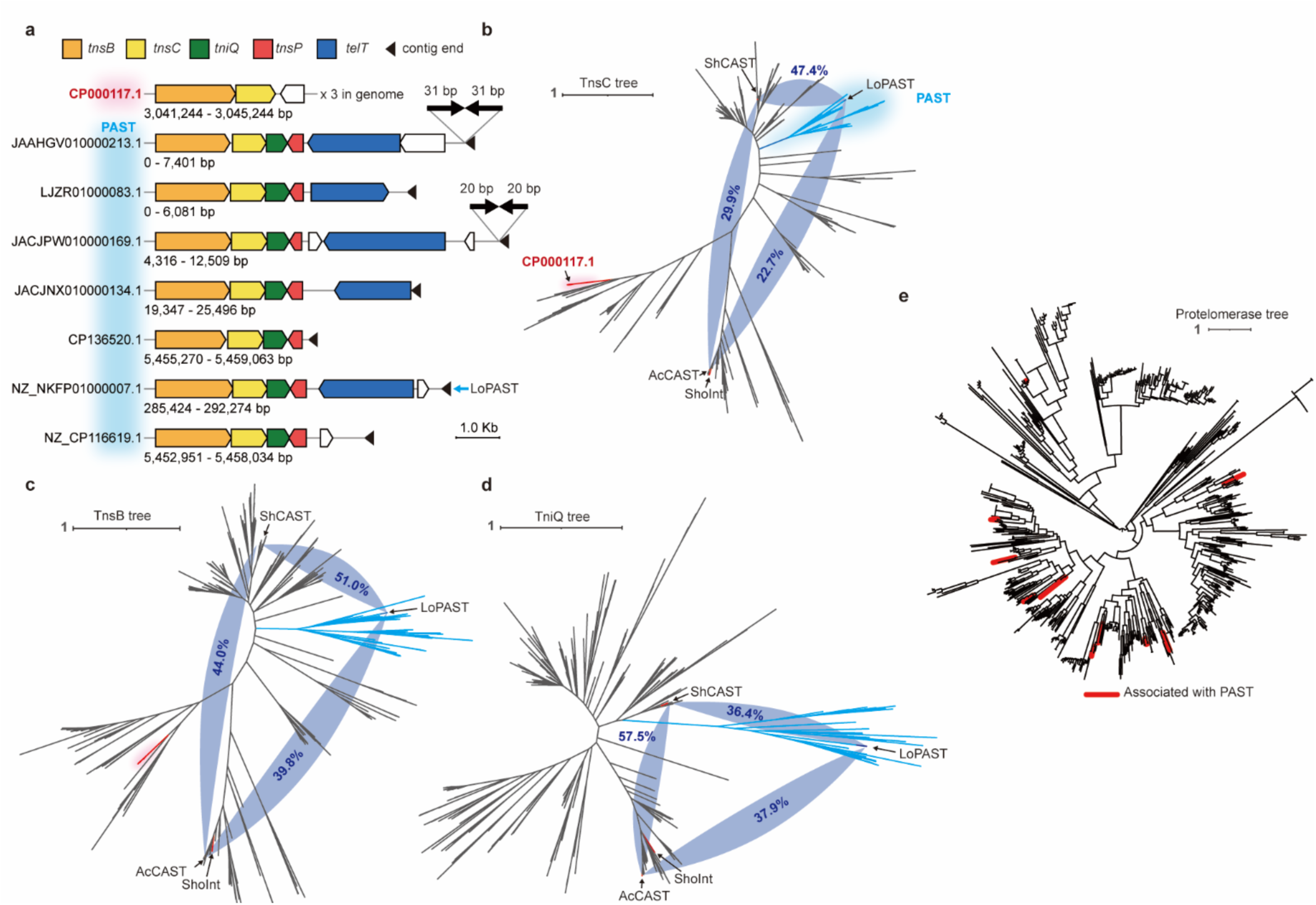
Protelomerase-associated transposons (PAST) related to CRISPR-Cas-associated transposons (CAST) form a distinct branch in cyanobacteria. (a) Various examples of PAST situated near the ends of contigs and a TnsBC non-PAST element which has 3 identical copies in the genome (CP000117.1 shown in red). *telT*: protelomerase, *tnsP*: small or found in PAST elements. (b,c,d) Similarity trees of TnsC, TnsB, and TniQ proteins from cyanobacterial Tn7-like transposons lacking TnsA reveal that PAST transposons (blue) and a group of TnsBC non-PAST transposons CP000117.1 are closely related to type V-K CASTs. The validated type V-K CASTs and the LoPAST tested in this study are labeled. The protein identities between the homologs of the labeled transposons are marked in the tapered dark-blue connections. (ShoInt is not used for comparison because it is a close homolog of AcCAST.) (e) Protein similarity tree of all putative protelomerases in cyanobacteria. The protelomerases associated with PASTs are marked as thickened red branches. PAST protelomerases do not form a single clade. The tree is midpoint rooted.

### Protelomerase-associated transposons (PAST) form a distinct group

Phylogenetic analysis of the 27 protelomerase-associated transposons (PAST) indicates that they are a new type of transposon that is distinct from the type V-K CAST elements. In all TnsB, TnsC, and TniQ similarity trees, PAST elements form a single clade (Fig. 1b,c,d), supporting that they belong to a distinguished group of transposons. The TnsB and TnsC of PASTs are closer to type V-K CASTs, while their TniQ is more diverged from type V-K CAST, implying a functional adaptation shifting away from Cas12k-guided transposition.

The putative protelomerase enzymes encoded in the PAST elements share little homology and have different domain architectures. Unlike TnsBCQ, the transposon-associated protelomerases do not form a single clade but scatter among all putative cyanobacterial protelomerases on the protein similarity tree (Fig. 1e). The lack of codivergence between TnsBCQ and protelomerase suggests that new protelomerase genes and their cognate ends are regularly captured by these transposons and are unlikely to be specifically adapted to the PAST elements.

### The transposon-associated protelomerase resolves PAST-born telomeres

A PAST element from *Leptolyngbya ohadii* IS1 (LoPAST) was selected to test our hypothesis on transposition activity and protelomerase co-option. LoPAST has two copies at different positions in the genome, suggesting that the transposon is active (Fig. 2a). Protelomerases are well-studied enzymes that form a hairpin at a cognate sequence they recognize^1,11^. While the activity of these enzymes is well-established, we wanted to confirm that the transposon-encoded examples can carry out the same function using the telomere sequence associated with the transposon. Although the sequences of the two LoPAST elements did not reveal the cognate telomere sequence, this sequence could be deduced from a close homolog in the same genome (79% amino acid identity). We expressed and purified the protelomerase (TelT) and incubated it with inverted repeat DNA that mimics the fused telomeres after replication. This proved that the LoPAST-encoded TelT can resolve the DNA into two hairpin fragments (Fig. 2b-c). Biochemical experiments with varying substrate lengths showed that TelT activity follows canonical chemistry and requires a substrate that minimally has 28bp inverted repeat sites, each predicated to be bound by protelomerase, consistent with previous protelomerase-DNA complex structures^11–14^ and AlphaFold prediction of a TelT-DNA complex (Fig. S1a-e). These experiments confirm protelomerase activity on PAST-born telomeres and supports our hypothesis that the PASTs are one-end transposons with a DNA hairpin on the other end.

**Fig. 2.**
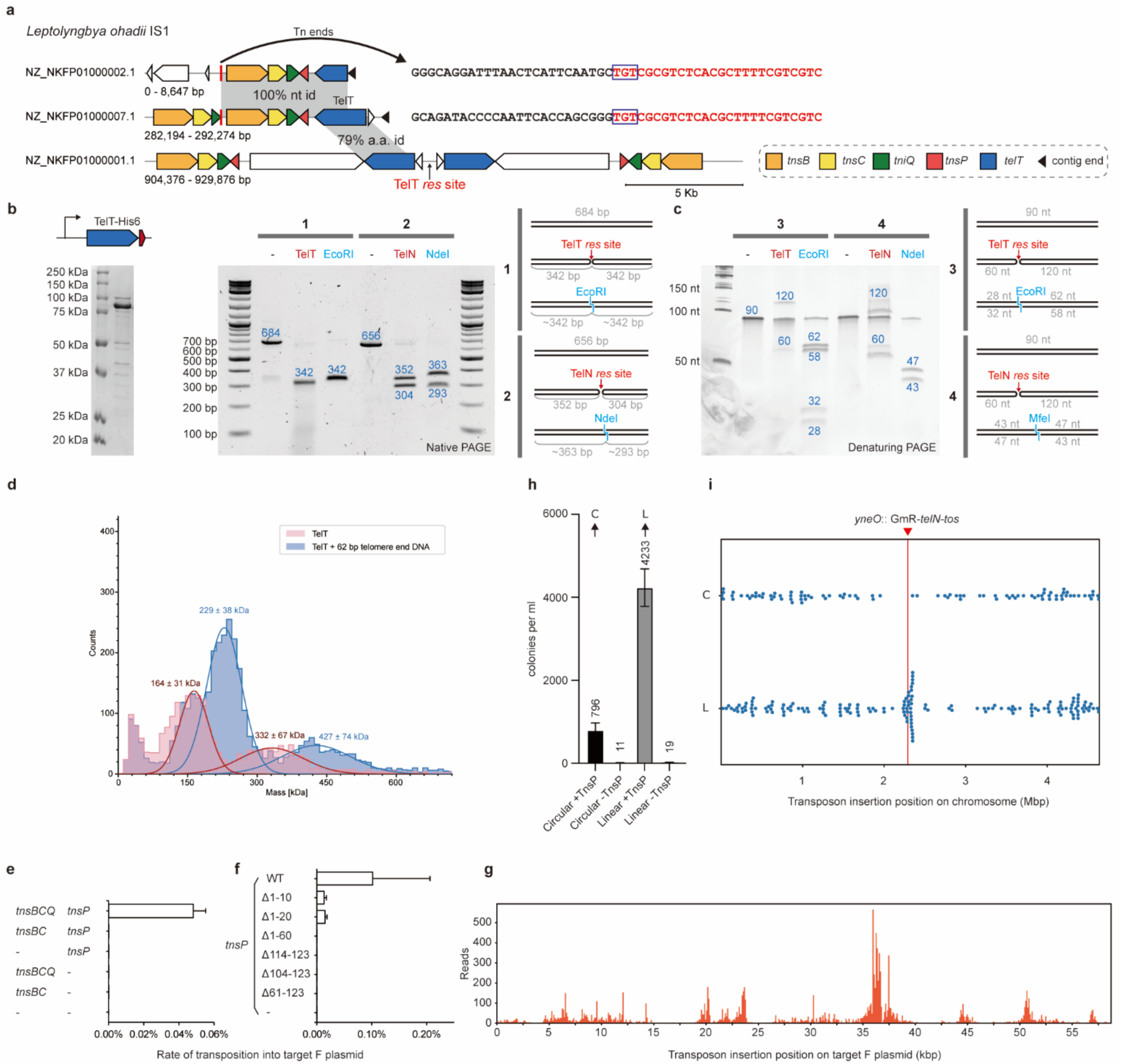
TelT from LoPAST functions as a protelomerase and LoPAST can transpose with two ends. (a) Identical copies of the LoPAST element found at different telomeres in the same genome (on the left) with different flanking sequence next to the “TGT” end of the element (on the right) suggests recent LoPAST transposition in the native host. A similar element is shown, likely captured in the DNA sequence after chromosomal replication and before resolution, forming a mirror image of the element. (b) Purified LoPAST protelomerase TelT (81.7 kDa, left) can resolve a 684 bp inverted repeat substrate with PAST-born telomere sequence into two 342 bp fragments (right). N15 phage protelomerase TelN and its resolution site *tos* and restriction enzyme digests are used as positive controls. (c) Denaturing PAGE shows hairpin formation upon TelT activity. 90 bp DNA duplexes are resolved into 60 nt and 120 nt ssDNA products, respectively (right). (d) Mass photometry analysis of TelT (81.7 kDa) in the absence (red) and presence of DNA (blue). TelT predominantly forms dimers (164 kDa). With telomere end DNA (38 kDa), a larger assembly is formed (229 kDa), representing a dimeric protein-DNA complex. (e) Results of the mating-out transposition assay with LoPAST. Transposition rates with various combinations of transposon genes show transposition into the target F plasmid only when TnsBCQ and TnsP are expressed. (f) Truncations of TnsP greatly reduce or eliminate its ability to stimulate transpositions. (g) Distribution of PAST insertion sites on the F-plasmid target mapped with DNA sequencing. (h-i) Transposition into the *E. coli* genome is found when TnsP is coexpressed with TnsBCQ. The linearized *E. coli* chromosome accumulated more transposition events proximal to the linear ends, suggesting LoPAST has the ability to target telomeres. (C: circular chromosome target, L: linear chromosome target.)

### Two transposon ends can support PAST transposition, contingent on TnsP

DNA transposons require two transposon ends that form an obligatory synapse with the transposase in the process of mobilization^11^. The close homology between transposition proteins encoded in the PAST and CAST elements implies that transposition of PAST elements should not be drastically different from type V-K CAST elements and DNA transposons in general. PASTs should still need two ends for transposition despite residing in genomes with only one transposon end. We hypothesized that PASTs could mobilize as an element with two ends, a configuration that is transiently found when DNA replication across the hairpin ends at the telomeres creates two inversely repeated transposon copies at the junction of the nascent sister chromosomes (Fig. S2). Non-covalent pairing of chromosome telomeres by protelomerase could also provide an opportunity for two separate one-ended transposons to be bridged, forming a substrate that was functionally a two-ended transposon. Previous work with protelomerase suggests diversity in the behavior of these enzymes. In the case of TelK, the protein does not associate with the hairpin products after resolution, while the TelA dimer can remain bound forming a linkage between the telomere ends ^12,13,15^. Similar to TelA, *in vitro* mass photometry and DNA binding assays demonstrated that the LoPAST TelT can mediate stable dimerization of resolved telomeres (Fig. 2d and Fig S1f).

A transposition assay in *E. coli* confirmed LoPAST activity as an element with two transposon ends (Fig. 2e)^16^. A mini-LoPAST transposon with two transposon ends flanking a kanamycin resistance gene was constructed into a plasmid to be used as a donor. We also tested the impact of a putative small open reading frame found in all PAST elements encoded downstream of the TnsBCQ encoding operon. When all LoPAST transposase genes TnsBCQ and the small open reading frame were expressed, transposition was readily detected, confirming that a configuration with two ends can support transposition. Truncation experiments with small open reading frame revealed that even a deletion of just 10 amino acids on the N- or C-terminus substantially diminished transposition (Fig. 2f). Mapping the insertions on the F plasmid target showed that the insertions are mainly non-specific (Fig. 2g). We refer to the small open reading frame as TnsP because it is conserved in the PAST elements and required for transposition with LoPAST.

We also analyzed PAST transposition into the *E. coli* chromosome to better understand target site selection. For this, the donor element with two ends was placed on a plasmid that could be easily cured from cells following a period of induction of the transposition functions. We also constructed a derivative of *E. coli* with a linear chromosome with hairpin ends to determine any underlying targeting features of the LoPAST element^17^. As with transposition into the F plasmid, TnsP was essential for transposition, and we reproducibly observed higher levels of transposition with a linear *E. coli* genome (Fig. 2h). Mapping transposition events with whole genome sequencing revealed a random component to transposition, however, insertions accumulated at a pronounced hotspot for transposon insertion near the telomeres in the linear *E. coli* chromosome (Fig. 2i).

### TnsP is a DNA binding protein that binds DNA without sequence specificity

TnsP has no homologs of known function, but structure prediction indicated that it forms homodimers encircling the DNA (see below). EMSA experiments indicated that TnsP binds short double-stranded DNA substrates with or without a hairpin end, but not single-stranded DNA (Figure S1g). TnsP shows no obvious bias for binding the telomere sequence within the LoPAST element, or a preference for DNA of a specific GC content, and binding is not affected by the purification tag (Fig S1h). The predicted ability of TnsP homodimers to encircle DNA is reminiscent of the behavior of the bacterial chromosome structuring protein GapR, which senses the super helicity of DNA and acts as an accessory factor to stimulate Type II topoisomerases^18^. An intriguing idea is that TnsP may interact with TniQ allowing it to sense features found at telomeres (see discussion), however, more research is needed to understand the specific mechanisms of transposition targeting to subtelomeric chromosome regions.

### Discovery of two additional families of telomeric transposons in *Streptomyces*

Linear plasmids and chromosomes are frequently found in actinobacteria, particularly in certain lineages such as *Rhodococcus*, *Kitasatospora*, and *Streptomyces*. Nearly all *Streptomyces* have linear chromosomes, and they tend to be genetically unstable, with mobile genetic elements enriched at their sub-telomeric regions^2^. Following the discovery of cyanobacterial telomeric transposons, we analyzed *Streptomyces* genomes in search of potential telomeric transposons.

Telomeric transposons are expected to be located exclusively near telomeres. We reasoned that if PAST elements exist as one-ended DNA transposons, related telomeric transposons should likely have a conserved gene orientation relative to the telomere. We screened and annotated complete *Streptomyces* chromosomes for DDE transposases (which includes TnsB), made a protein similarity tree, and plotted their relative locations and orientations. The analysis showed that the distribution of transposases differs across families, but they are generally enriched near the ends of chromosomes (Fig. S3). However, two transposon families showed a conserved gene orientation and were found essentially exclusively at the sub telomeric region of chromosomes, one belonging to TnsBC transposons with a transposase like TnsB from Tn7 and TnsC but lacks the TnsA and TnsD/TniQ proteins normally associated with Tn7-like transposons and another belonging to Tn7-like transposons (Fig, S3).

### Telomeric Tn7-like transposons in *Streptomyces* are associated with telomere-maintaining genes and the putative conjugation helicase TtrA

To characterize Tn7-like transposon distribution in these species, we conducted a computational search for Tn7-like elements in annotated *Streptomyces* genomes available from the NCBI database. Approximately 40% of annotated *Streptomyces* genomes carry at least one putative Tn7-like transposon, and about 40% of these belong to the branch of putative telomeric transposons (∼16% genomes containing at least one Tn7-like telomeric transposon)(Fig. 3a).

**Fig. 3.**
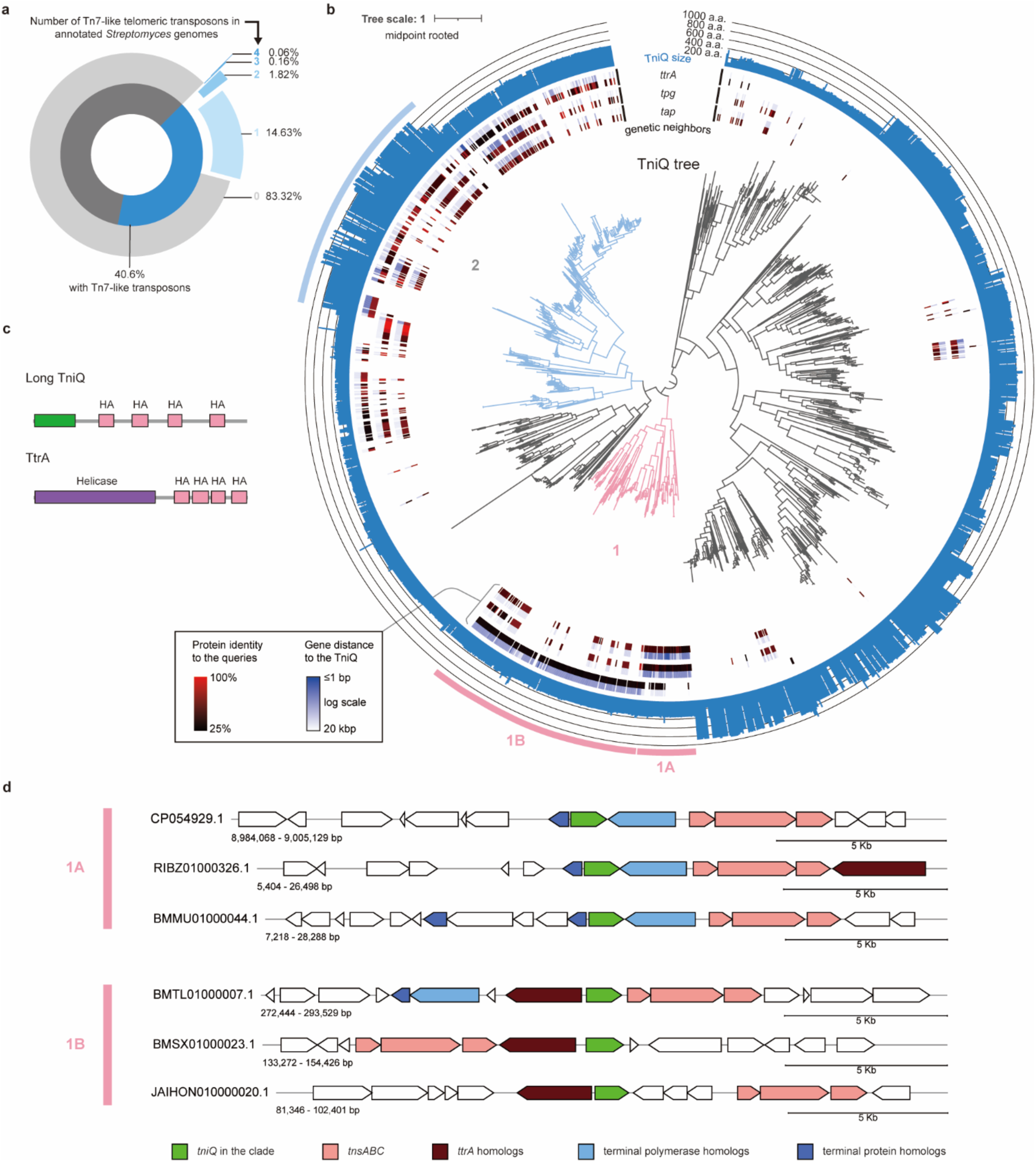
Two independently evolved families of telomeric transposons are found in *Streptomyces* genomes. (a) Breakdown of Tn7-like elements found in *Streptomyces* genomes. Tn7-like transposons appear in 40% of annotated *Streptomyces* genomes (dark blue), with over 16% containing one or more Tn7-like telomeric elements (light blue). (b) TniQ tree of Tn7-like transposons in *Streptomyces*. The inner rings show genetic association with homologs of terminal polymerase (*tap*), terminal protein (*tpg*), and conjugation helicase (*ttrA*) each indicated with two color bands. One color band shows the distance between the homolog and the TniQ; the other shows the protein identity between the homolog and query as indicated in the heatmaps.The outer ring indicates the amino acid (a.a.) length of the TniQ in the tree as a histogram. The two branches of telomeric TniQ are colored pink (clade 1) and light blue (clade 2). (c) Large TniQ of clade 2 telomeric transposons have tandem HA (helicase-associated) domains similar to TtrA homologs. (d) Clade 1 TniQ has a strong genetic association with *tap*, *tpg*, and *ttrA*. Clade 1A *tniQ* are sandwiched by *tpg* and *tap*; clade 1B *tniQ* always have *ttrA* upstream in the opposite orientation.

Notably, we observed that many Tn7-like telomeric transposons are closely associated with genes responsible for maintaining telomeres. This was analogous to the finding that PAST elements from cyanobacteria maintain their own protelomerase and cognate telomere end sequence. Unlike cyanobacterial telomeres, the telomeres of linear chromosomes in *Streptomyces* are maintained by terminal proteins, which prime DNA replication and remain covalently bound to the DNA at the telomere end. A specialized B-family polymerase catalyzes DNA replication, and its gene is commonly found forming an operon with the terminal protein gene. Across *Streptomyces* there are three characterized terminal polymerase and terminal protein pairs, Tap-Tpg, Tac-Tpc, and GtpB-GtpA^19–22^, which share no sequence homology.

In addition to the telomere maintenance proteins, we found homologs of the conjugation helicase gene *ttrA* to be associated with telomeric transposons. The *ttrA* gene product is known to be located in the sub telomeric region and involved in the transfer of telomeric ends between cells^23^ (Fig. S4 and S5). We investigated the relationship between the Tn7-like transposition genes and the genes encoding the TtrA conjugation helicase and Tap-Tpg, Tac-Tpc, and GtpB-GtpA in *Streptomyces* genomes. Mapping the genetic distance between the genes associated with telomere function and TniQ revealed a close association in the telomeric transposons. (Fig. 3, S6). Tn7-like telomeric transposons can be subdivided into two clades based on TniQ protein similarity (Fig. 3). Notably, clade 1 Tn7-like telomeric transposons displayed a particularly robust genetic connection in this regard. The clade 1A TniQ is sandwiched between terminal polymerase and terminal protein genes, ensuring the genetic linkage (Fig. 3d, S4); clade 1B TniQ is always associated with the *ttrA* gene (Fig. 3d, S5). As expected, the *ttrA* gene are also found at telomeres in *Streptomyces* genomes (Fig S7). The genetic association further supports that these transposons are adapted to a telomeric lifestyle.

While transposons generally accumulate at the ends of chromosomes, we found evidence of telomere-targeting in some *Streptomyces* genomes. For example, a Tn7-like telomeric transposon is found at three telomeres on the chromosome and a megaplasmid in *Streptomyces* CLI2509 (Fig. S8). These three copies of transposons are flanked on one side by TGT/ACA, the typical end sequence of Tn7-like transposons, supporting the idea that these copies are mobilized by transposition. Movement of telomeric transposons can also be observed between different *Streptomyces* species. For instance, near identical elements can be found in *Streptomyces coelicolor* A3(2) and *Streptomyces olivaceus* N11-26, and they are also flanked on one side by TGT/ACA (Fig. S8).

*Streptomyces* Tn7-like telomeric transposons display the orientation bias that aligns with the feature of one-end transposons and our investigation revealed only one putative transposon end. In *S. coelicolor* A3(2), the sequence space between the transposon end and the characterized telomere is densely populated with coding sequences, making it unlikely for us to overlook the presence of the other transposon end if it existed. Thus, akin to PAST, it seems that the borders of these telomeric transposons are delineated by one transposon end and one telomere.

### Loss of TnsA nuclease activity in *Streptomyces* Tn7-like telomeric transposons allows one-ended transposition

Similar to PAST, having only one transposon end poses multiple challenges for the life cycle of *Streptomyces* telomeric transposons. One challenge is how to associate two transposon ends for DNA transposition. In *Streptomyces*, this can possibly be mediated through non-covalent interaction between telomere maintenance proteins, which have been shown to be strong enough to constrain supercoiling between the DNAs where they associate^24^ and can likely bridge transposon ends suitably to allow transposition. A second issue that is specific to Tn7-like elements is the process of cut-and-paste transposition, which is facilitated by the TnsA nuclease. In regular Tn7-like transposons with two ends, TnsB cuts one DNA strand at each end of the element and strand transfer results in a Shapiro intermediate with one strand of the element connected to the insertion site in the target DNA and the other still connected at the donor DNA (Fig. 4)^25^. In prototypic Tn7 transposition, TnsA cuts the strand connecting the transposon to the donor DNA at each end of the element, resulting in cut-and-paste transposition and leaving double-strand breaks on the donor DNA^26^. Without TnsA cutting, replication will progress through the transposon, duplicate it, and result in the cointegration of donor DNA into target DNA (a process known as replicative transposition)^27^. However, if the two transposon ends are not covalently connected, as we hypothesized for *Streptomyces* telomeric transposons, replicative transposition will not result in cointegrate. The donor and target DNA can resolve themselves after DNA replication. In this scenario, TnsA activity will only cause unnecessary DSBs in the donor DNA, potentially leading to the loss of telomeric transposons and the telomeres themselves, which will likely lead to cell death. Therefore, we predicted that the TnsA protein of Tn7-like telomeric transposons might lose their native nuclease activity as a step in adapting to life as a one ended transposon.

**Fig 4.**
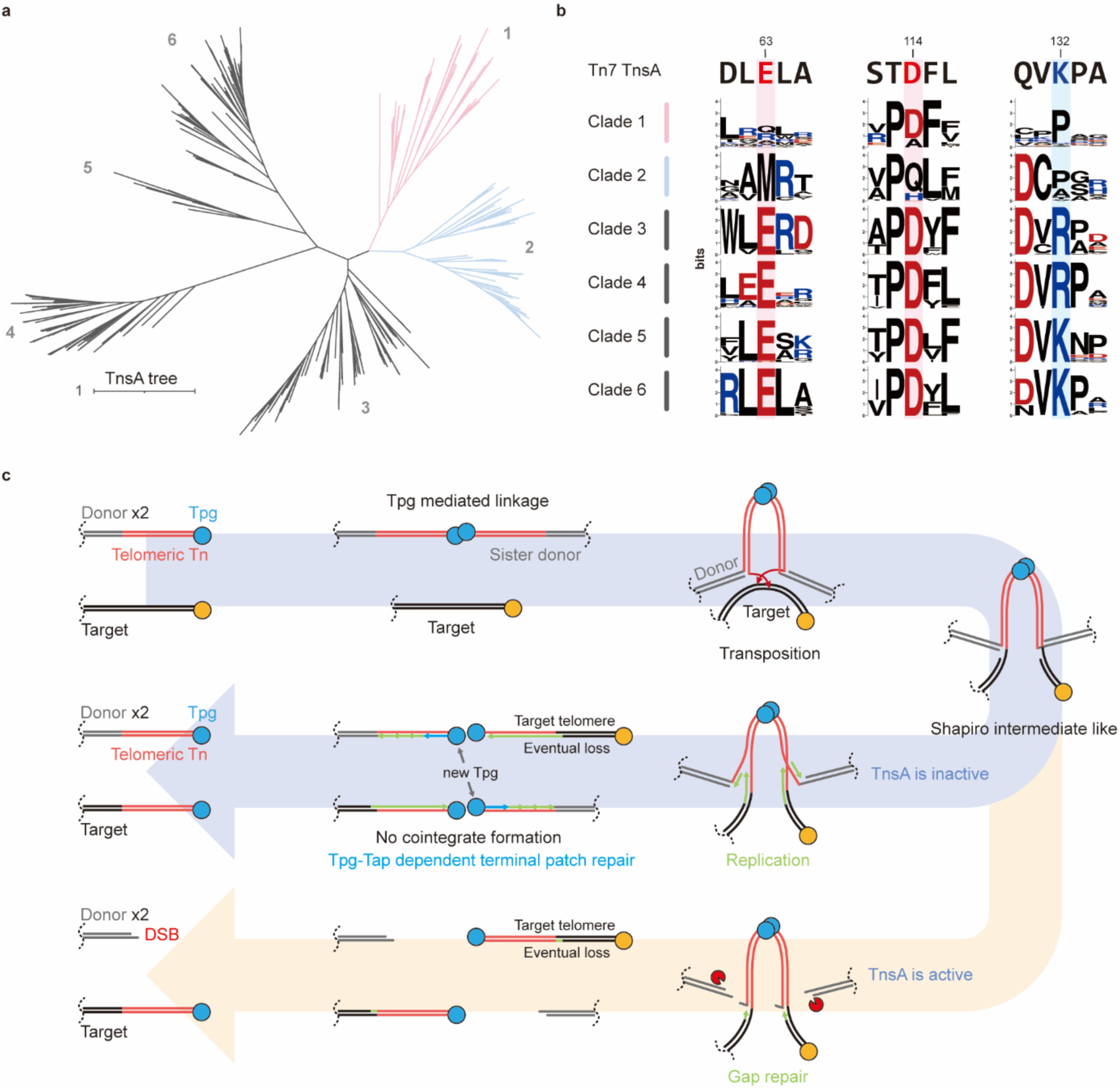
One branch of Tn7-like transposons from *Streptomyces* has TnsA proteins with a change in the active site residues with a known consequence for the transposition mechanism. (a) The TnsA tree of *Streptomyces* Tn7-like transposons with TnsA clades of Tn7-like telomeric transposons colored (TnsA proteins were deduplicated with a 95% cut off). (b) The amino sequence consensus near the active site residues of TnsA proteins of different clades. Conserved charged residues are needed for DNA cleavage. The active site residues of telomeric transposon TnsA deviate from conserved sequences. The active site sequences of Tn7 TnsA are shown for comparison. (c) Scheme of the putative transposition pathway of Tn7-like telomeric transposons in *Streptomyces*. Two telomeric transposons form a transpososome through the non-covalent interaction between terminal proteins. Following transposition, the donor and target DNA form a Shapiro intermediate-like structure. Without TnsA cutting, this structure can resolve itself after DNA replication, resulting in transposon copies both at the donor and target sites. In the hypothetical pathway shown in beige, TnsA cutting, would result in cut-and-paste transposition, leaving double-strand breaks on two donor DNAs. Therefore, in the case of telomeric transposons, TnsA cutting would be highly disadvantageous for the element and host.

To investigate this prediction, *Streptomyces* TnsA sequences were classified into six clades based on their similarity (Fig. 4a). The analysis revealed that the three active site residues crucial for executing DNA cleavage are conserved in non-telomeric transposons (Fig. 4b). In contrast, these residues exhibit alterations in both clades of *Streptomyces* telomeric transposons, aligning precisely with our prediction. Despite losing the nuclease activity, these telomeric transposons still retain TnsA. This suggests that the protein is essential for transposition even in its nuclease-dead state, resembling the archetypical Tn7 and I-F3 CAST^27,28^. Another difference found in the clade 2 elements, is that they contain two TnsB transposases, one of which lacks the required active site residue (Fig. S9).

### Gene context indicates diverse targeting or maintenance strategies in TnsBC telomeric transposons

Another family of telomeric transposons identified in *Streptomyces* is related to Tn7-like transposons but lacks *tnsA* or *tniQ* genes. For convenience, we call this transposon family TnsBC elements. All transposons within the family encode a transposase gene (TnsB) and at least one putative AAA+ regulator (TnsC) downstream. TnsBC telomeric transposons can be found in over 17% of annotated *Streptomyces* genomes. We also found evidence of recent transposition in *Streptomyces* sp. YIM121038 and *Streptomyces cadmiisoli* strain ZFG47, where the same TnsBC telomeric transposon populates chromosome telomeres and telomeres of a linear megaplasmid, with most transposon ends demarcated with TGT/ACA, as in Tn7-like transposons (Fig. S10).

Notably, we found that the one-ended transposon evolved from a typical two-ended transposon: elements in the basal branch of this family are not telomeric transposons and retain both transposon ends as is normally required for DNA transposons (Fig 5). This is like the association between one-ended LoPAST elements that branch separately from the V-K CAST elements with two transposon ends (Fig. 1). Basal branch TnsBC elements with two transposon ends are often located on plasmids near conjugation genes, and in some cases, both transposon ends and 5 bp target site duplication can be identified (“SR” in Fig. 5b). Another feature also supports that these elements use a replicative transposition mechanism: they appear to associate with a serine recombinase (“SR” in Fig. 5b). The serine recombinase likely functions as a resolvase for cointegrate resolution, something that is not required for one-end telomeric transposons according to our model.

**Figure 5.**
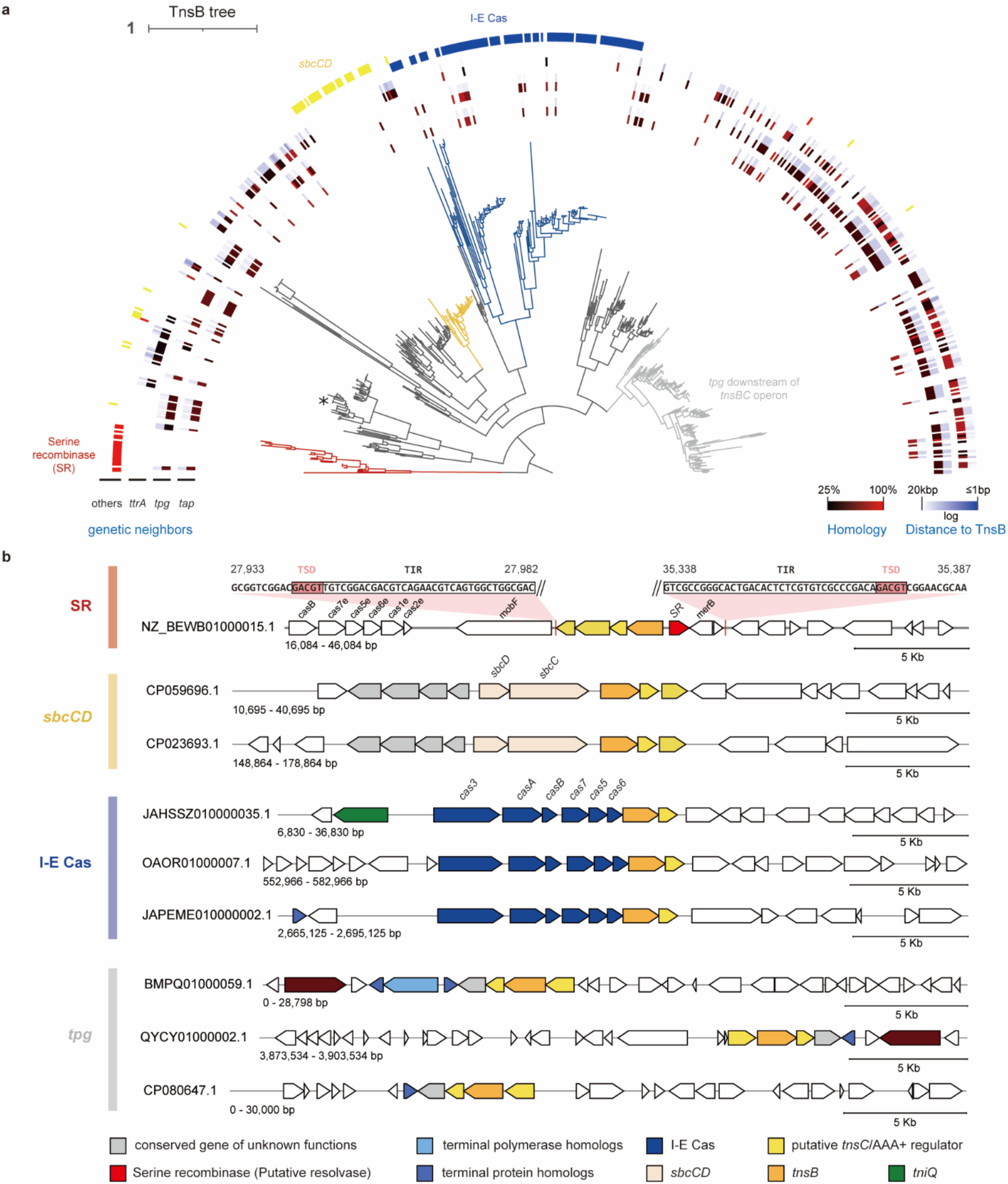
Diversity of TnsBC telomeric transposons in *Streptomyces* genomes. (a) The subtree contains the TnsBC telomeric transposons and their close relatives. The asterisk * marks the transposons shown in supplementary Figure S10. The tree is rooted with Tn7-like transposons in *Streptomyces*. Genetic association with homologs of terminal polymerase (*tap*), terminal protein (*tpg*), and conjugation helicase (*ttrA*) are displayed as colored rings. Other conserved associated genes (within 20 kbp) are displayed as colored rings. (b) Examples of selected gene configurations found in the subbranches. Elements in the basal branch (“SR”) have two transposon ends, target site duplications, and carry a serine recombinase gene, consistent with canonical replicative DNA transposition. Elements from other subbranches associate with telomer maintenance, DNA repair and targeting functions.

Each of the TnsBC telomeric transposon groups we identified are associated with specific accessory genes. One subbranch is associated with *sbcCD* genes, encoding the well-studied DNA repair nuclease, and a four gene operon, including proteins predicated to have exonuclease and helicase activities (“*sbcCD*” in Fig. 5). This suggests a lifestyle that may involve double strand break DNA damage and repair. Another subbranch is associated with terminal protein genes (“*tpg*” in Fig. 5). It has a *tpg* homolog downstream of the *tnsBC* operon, facsimile to the Tn7-like telomeric transposons, supporting that these transposons are also adapted to telomere-targeting lifestyle. Unexpectedly, one subbranch is linked to type I-E CRISPR system, with its *tnsBC* genes integrated into the type I-E cascade operon (“I-E Cas” in Fig. 5). While these elements were previously described and initially hypothesized to have TnsBC substituting Cas1-Cas2 for spacer acquisition^7^, our study reveals that these genetic elements are, in fact, type I-E CRISPR-associated telomeric transposons (see below).

### Type I-E CAST are one-ended telomeric transposons that move with two transposon ends

Canonical type I-E CRISPR systems in *Streptomyces* feature the *cas3* gene, cascade operon with *cas1* and *cas2* at the end, and long CRISPR arrays flanking the Cas genes. In contrast, type I-E CAST incorporates the *tnsBC* genes within the operon instead of *cas1-2*, and its CRISPR arrays are exclusively situated upstream of *cas3*. The Cas proteins of type I-E CAST constitute a separate clade distinct from other conventional type I-E Cas on protein similarity trees (Fig. 6a), suggesting a stable genetic association and coevolution with the transposition genes. We examined the positions and orientations of type I-E Cascade operons on assembled *Streptomyces* chromosomes, which revealed that only the type I-E CAST consistently resides in proximity to the telomere with a conserved orientation (Fig. 6b). To assess whether the type I-E CRISPR system associated with the *tnsBC* is adapted for telomere targeting, we analyzed spacers from both canonical and type I-E CAST arrays and compared their protospacer compositions. Among the more than 29,000 non-redundant spacers from canonical type I-E CRISPR, we found a proportion to target coding sequences annotated as phage (4.43%) or plasmid conjugation (1.03%) related, and only about 0.3% of these spacers matched telomere-related genes (*ttrA*, *tap*, *tpg*) (Fig. 6c). Conversely, of the 674 non-redundant spacers from telomeric type I-E CAST, a much higher proportion targets telomere-associated *ttrA* (5.93%), and only 1% hits genes annotated as phage or plasmid conjugation-related (Fig. 6d). Interestingly, numerous spacers match *tnsB* or *tnsC* genes (5.64%), all of which belong to either Tn7-like or TnsBC telomeric transposons. In total, 13.5% of non-redundant spacers from type I-E CAST targets telomere associated genes. Furthermore, excluding phage, plasmid conjugation, and known telomere associated genes, the protospacers of type I-E CAST are predominantly located near the telomere on assembled *Streptomyces* chromosomes. While the canonical type I-E CRISPR also exhibits a modest enrichment of protospacers near telomeres, it is not nearly as pronounced as in type I-E CAST (Fig 6b). The protospacer compositions strongly indicate that the type I-E CRISPR system that is associated with transposons is specifically adapted for telomere targeting.

**Fig. 6.**
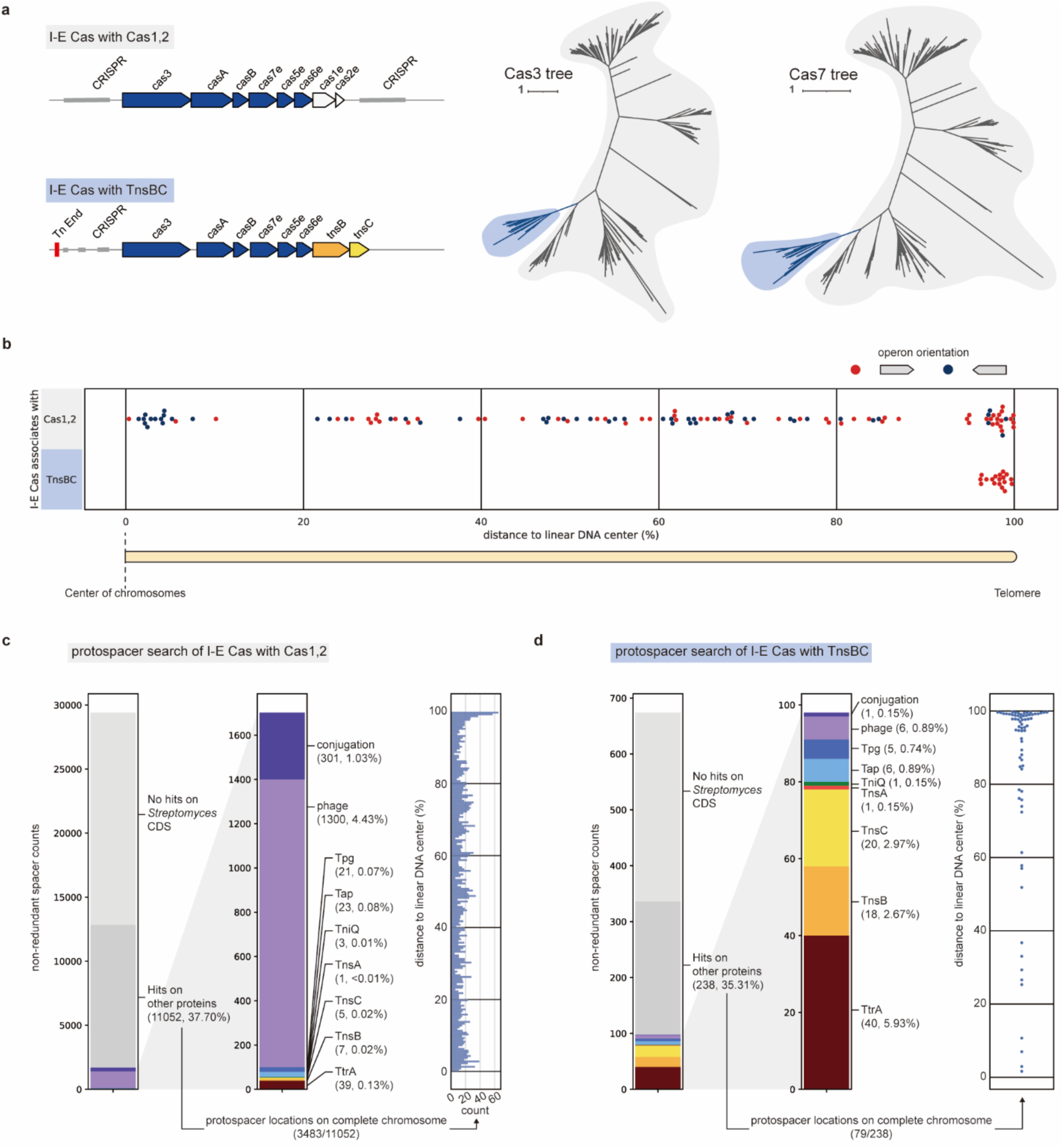
Analysis of the type I-E CRISPR-associated transposons (I-E CAST). (a) Genetic organization of typical type I-E system associated with Cas1-2 and I-E CAST. The Cas proteins of I-E CAST form a single clade. Only Cas3 and Cas7 trees are shown here, but other Cas proteins consistently showed a single clade for the I-E CAST components. (b) The location of type I-E systems associated with *cas1-2* or *tnsBC* genes on complete *Streptomyces* chromosomes plotted according to their relative positions to the center of linear chromosomes. The orientations of operons are indicated in red or black. (c) left and middle: A protospacer search of type I-E systems associated with Cas1-2. Many spacers match phage and conjugation genes, with few matches to known telomere-related genes. (d) left and middle: A protospacer search of type I-E system associated with *tnsBC*. Most spacer matches are to known telomere-related genes and telomeric transposon genes, and relatively few matches to phage and conjugation genes. (c-d right) Chromosomal locations of protospacers that are not known to be telomere-related or phage, conjugation, or telomeric transposon-related. The protospacers of type I-E CAST are enriched near telomeres, which is not found with canonical type I-E systems.

In each of the cases above, telomeric transposons reside in the host genome with one traditional transposon end and a telomere flanking the transposon and cargo genes. While this observation suggests a completely novel mechanism may be used for transposition, the LoPAST element was found to move like a DNA transposon with two ends. We set up a system to test a putative I-E CAST element in *E. coli* but were unable to detect transposition when expressing the wild type TnsB and TnsC proteins in absence of targeting components (Fig S11a). Previously it was shown with prototypic Tn7, that inactivating the Walker B motif of the TnsC regulator protein allowed a gain-of-function phenotype, where transposition could occur without the target site selection systems (TnsD or TnsE)^29,30^. We tested the analogous mutations in the I-E CAST system and observed detectable transposition with TnsC(E134A) and TnsC(E134K), as well as robust transposition levels with the TnsC(E134Q) change (Fig. S11b, like the finding with prototypic Tn7^30^. DNA sequencing confirmed that transposition produced the predicted target site duplication (Fig. S11b). Although we did not recapitulate transposon targeting with these experiments in *E. coli*, we were able to show transposition supporting the idea that it is a bone fide I-E CAST transposon that moves using two transposon ends.

### Insights into the Transposition Mechanism of Type I-E CAST

All previously characterized CASTs belong to Tn7-like transposons, displaying similar adaptations despite evolving independently. The CRISPR systems co-opted by CASTs have undergone the loss of *cas1*-*cas2* for spacer acquisition, and they either lost genes for interference or have inactivating mutations at the nuclease, leaving only CRISPR-Cascade for target recognition, which is interfaced with the TniQ protein to guide transposition. Type I-E CAST is not a Tn7-like transposon, rather a TnsBC element. It lacks TniQ and must interact with the co-opted CRISPR system differently than previously characterized systems. Although it lacks *cas1*-*cas2*, it retains *cas3* with an intact nuclease active site. Additionally, we cannot identify protospacers adjacent to the predicted transposon ends, something that is inconsistent with the mechanism of guide-RNA directed transposition as found with the other CASTs. These observations collectively suggest a distinct relationship between the transposition and CRISPR-Cas systems.

To gain insights into how the type I-E CASTs might use the co-opted CRISPR system to assist transposition, we searched for DNA sequences that were homologous to where the transposon insertion resided but without an insertion (pre-insertion sites). Most pre-insertion homologs are short, likely due to frequent rearrangement of subtelomeric regions, or lack of sequenced representatives. Nonetheless, we found two cases of long pre-insertion homologs (match >20 kbp upstream of transposon ends) with high sequence identity (>99%) (Fig. S11a). Both sequences that are homologous to chromosomes in other bacteria that contain a I-E CAST insertion provide evidence of the relationship between CRISPR-Cas-mediated degradation and the resulting transposition events. In both cases a protospacer that matches the spacer found in the type I-E CAST element in the homologous sequences is proximal to the insertion site (83 bp and 339 bp apart, respectively). The protospacers are facing away from telomeres, the more common orientation of type I-E CAST protospacers (77 %)(Fig. S11c); and based on this configuration they must have been deleted during the process of recognition and integration (Fig. S11a). While further investigation is required to elucidate the precise targeting mechanism of type I-E CAST transposition, these clues provide a preliminary understanding of certain aspects of the process. The process likely starts with type I-E Cascade targeting subtelomeric genes with the protospacers oriented away from telomeres (Fig. S12). With this orientation Cas3 is activated to unidirectionally degrade DNA toward telomeres. In the model, transposition is then guided to the adjacent region of protospacers through the interaction of TnsBC with cascade or DNA repair intermediates. We suggest that an interaction between TnsBC with DNA repair intermediates is more likely to be the case as the varying distances between protospacers and insertion positions contradict the expectation of site-specific recruitment via the Cascade.

### *Streptomyces* telomeric transposons play a prevalent role in the turnover of *Streptomyces* telomeres

Tn7-like/TnsBC telomeric transposons are identified in over 31% of annotated *Streptomyces* genomes. These transposons exhibit mobility between linear chromosomes and plasmids and disperse among various hosts, effectively functioning as mobile telomeres. The cargo sizes of telomeric transposons (approximated by distance from transposases to the ends of assembled chromosome) can range from almost none to over a hundred kilo base pairs (Fig. S13a). This means that telomeric transposons have the capacity to mobilize numerous genes within and between genomes, altering the telomere and subtelomere sequences. We also observed tandem insertions of telomeric transposons, sometimes disrupting an element that previously resided at the telomere, showing that interaction and competition between these abundant transposons may also lead to the diversification, hybridization, and evolution of both the transposons and the affected telomers/genomes (Fig. S13).

## Discussion

Convergent evolution is a prominent theme across mobile genetic elements. Different transposons emerged to employ similar transposition mechanisms, like the recursive evolution of Tn7-like CRISPR-guided transposons. These transposons occupy similar genomic niches, such as the centromere and telomere targeting transposons in Eukaryotes, and tRNA-targeting mobile genetic elements in bacteria. The three families of independently evolved bacterial telomeric transposons found in this study further exemplify the phenomena of convergent evolution. Moreover, these telomeric transposons replace and take up the role of telomeres, in a way akin to *Drosophila* telomeric transposons, despite their fundamental differences in transposition and telomere maintenance mechanisms.

An ability for transposons to preferentially target bacterial telomeres has numerous advantages including the ability to avoid insertional inactivation of essential genes, which are consolidated more centrally in chromosomes^2^. In the case of *Streptomyces* and related bacteria, the telomeric region is also mobilized between bacteria in an incompletely understood conjugation process, promoting gene transfer and adaptation^31^. Interestingly, many telomeric transposons also encode a special helicase that is involved in conjugation, suggesting that the process of cell-to-cell transfer has been coopted by these transposons. Transposition of a one-end telomeric transposon results in the formation of a new telomere that is dependent on the telomere maintenance system encoded in the element, a strategy that can prevent the loss of the mobile element. Like native telomeres in cyanobacteria, the LoPAST TelT will make a hairpin structure that will solve the end-replication problem and not be recognized as a DNA double strand break. Similarly, the telomere maintenance systems associated with the telomeric transposons in *Streptomyces* will also provide these essential functions.

We note that transposition of a telomeric transposon randomly across the chromosome would be highly detrimental, leading to the loss of essential genes in large arms of the chromosome. Thus, telomeric transposons would benefit from a mechanism to specifically target telomeres and subtelomeric regions. Interestingly, we could show that the LoPAST transposition system was stimulated in *E. coli* strains where the chromosome was maintained as a linear molecule and transposition events accumulated at the subtelomeric region. However, the targeting mechanism remains to be established. An exciting yet speculative model would be that TnsP may act as a sensor of DNA super helicity as shown with an analogous protein GapR that encircles DNA and helps modulate the topoisomerase activity in some bacteria. Interestingly, structure predication suggests that like GapR, TnsP is predicated to encircle DNA with a positively charged surface inside the assembly that would be favorable for this interaction (Fig S14). Mechanistic details will require further investigation to understand the similarities between GapR and TnsP function in sensing the supercoiling state of DNA and the role this plays in target site selection.

TniQ generally mediates target selection in Tn7-like transposons and special adaptations are likely present in the telomeric transposons identified in this work. In LoPAST, TniQ relies on TnsP for transposition, suggesting it may act similar to the function of TnsF in Tn6022 elements. In most Tn7-like elements, the TniQ domain is covalently connected to one of many types of DNA binding domains that provide sequence specificity for target site selection. The TniQ from Tn6022 forms a non-covalent interaction with the TnsF DNA binding protein to target transposition to the specific DNA sequence that is recognized by this transposon^32,33^. Consistent with this idea, structure prediction for the TnsC/TniQ/TnsP complex on DNA using AlphaFold3 indicated a specific interaction between TniQ and TnsP, possibly allowing TnsP-dependent recruitment of the TnsBCQ machinery (Fig S15). In *Streptomyces*, some TniQ proteins in the clade 2 Tn7-like telomeric transposons have tandem Helicase Associated (HA) domains homologous to the domains in the telomere recognition protein TtrA (Fig. 2c). HA domains are named because they are found in multiple copies in some bacterial helicase proteins and may play a role in binding to DNA^34^. Further study will be needed to know if they recognize features/sequences associated with telomeres. The *Streptomyces* telomeres are genetically unstable and susceptible to DNA breakage. It is possible that features of DNA double-strand breaks, or the associated repair processes are targets of various TnsBC telomeric transposons. This is supported by the genetic association between one TnsBC transposon subbranch with *sbcCD*, whose homologous proteins are known to process broken DNA ends. The association with a special subbranch of type I-E CRISPR-Cas systems provides another link with DNA double strand breaks (see below).

Our work also clarifies the basic functioning of a subbranch of type I-E CRISPR-Cas systems that was found to associate with genes encoding TnsB and TnsC transposition proteins. These systems encode a Cas3 helicase/nuclease but lack a canonical Cas1Cas2 spacer acquisition system. Initial work with this system did not identify transposon ends flanking the elements, and the suggestion was made that the TnsBTnsC proteins might replace the spacer acquisition function of Cas1Cas2^7^. Here we show that these systems are I-E CRISPR-associated transposons (CAST) with one standard transposon end and one end that is a telomere, explaining the initial inability to identify the expected features on both sides of the element. Like for the LoPAST element, we can show *Streptomyces* telomeric transposon uses two ends for transposition as expected for DNA transposons. Activating transposition with the I-E CAST system in the heterologous *E. coli* host required bypassing the normal target site selection system using a TnsC mutant allele, a strategy inspired by work with the prototypic Tn7 system^29,30^. Additional research will be needed to understand the specific mechanism of target identification in the type I-E CAST systems. Type I-E CAST have spacers that specifically recognize sequences enriched at subtelomeric regions suggesting that homologous recombination could also function to mobilize these elements as suggested for the selfish spread of TnpB-encoding elements^35^. Type I-E CAST are also enriched in spacers that target other telomeric transposons, which would facilitate competition between telomeric elements.

This work reveals telomeric transposons as important players for genome maintenance and diversification in *Streptomyces* and related bacteria. Telomeric transposons are found in ∼30% of *Streptomyces* genomes. Interestingly, we find that even the archetypical telomere from the model organism *Streptomyces coelicolor* A3(2) is carried by a Tn7-like telomeric transposon. Telomeric transposons are not static inhabitants of the genomes where they reside. Although it would be challenging to quantify given the quality of available genome sequences, we have identified numerous instances where successive telomere turnover was facilitated by telomeric transposons. Homologous DNA recombination associated with the repair of DNA double-strand breaks will also provide another mechanism for allowing telomeric transposons to spread within and between individual bacteria and diversify genomic termini. In this way, *Streptomyces* genomes pose as battlegrounds for telomeric transposons and the cargo they carry.

## Methods

### Bioinformatic discovery of Tn7-like and TnsBC transposons in *Streptomyces* and *Cyanobacteria*

Annotated *Streptomyces* and cyanobacteria genomes were downloaded from NCBI. In total, there were 3,130 *Streptomyces* genomes and 2,894 cyanobacteria genomes. Tn7-like/TnsBC transposons were identified by scanning genomes with profile HMMs associated with TnsA (PF08722, PF08721, NF033179), TnsB (PF00665, PF09299), TnsC (PF11426, PF05621, PF13401), TniQ (PF06527, PF15978) downloaded from the European Bioinformatics Institute (EMBL-EBI) Pfam database and NCBI, utilizing the hmmsearch tool of HMMER v3.4^36^. Operonized TnsBC/TnsABC are assembled based on their proximity and orientation, adjacent TniQ are considered members of the same transposons with the TnsBC/TnsABC.

### Bacterial strains and growth conditions

Protein expression for protein purification was done in *E. coli* strain BL21-AI (TelT) or BL21 (TnsP). All in vivo transposition assays were done in *E. coli* strain background BW27783. Bacteria were grown in LB, TB or M9 media as indicated in the individual experiments^37^. Antibiotics were used at the following concentrations to maintain plasmids (Supplementary Table 1) and select for exconjugants or transposition events in the individual experiments, 100 μg/ml carbenicillin, 30 μg/ml chloramphenicol, 10 μg/ml gentamycin, 50 μg/ml kanamycin, 20 μg/ml nalidixic acid, 100 μg/ml rifampicin, 50 μg/ml spectinomycin.

### Protein expression and purification

A vector designed to express the C-terminally His_6_-tagged LoPAST protelomerase (TelT-His_6_) gene from a T7 promoter was synthesized by Twist Bioscience (pET29b backbone, Supplementary Table 1). *E. coli* BL21-AI cells were transformed with the TelT-His_6_ expression plasmid, colony purified and cultured overnight in 5 ml LB with 100 μg/ml ampicillin at 37°C. Cells were diluted into 50 ml LB the next day and inoculated into 500 ml TB medium upon reaching the log phase. The 500 ml TB culture was agitated in a shaker at 37°C until an optical density of 0.5 (O.D. 600 nm) at which point the temperature of the culture was gradually decreased to 16°C. Protein expression was induced with 0.1 mM IPTG and 0.2% arabinose when the O.D. 600 reached 0.6. After 16 hours of expression, the culture was spun down for protein purification. Cells were resuspended in lysis buffer (500 mM NaCl, 25 mM HEPES pH 7.5, 10% glycerol, 5 mM DTT) supplemented with 1mM PMSF and sonicated for lysis. Following sonication, cell debris was pelleted by centrifugation at 72,000 g for 2 hours at 4 °C, and the supernatant was collected. Imidazole was added to the supernatant to a final concentration of 20 mM and applied to a 2 mL Ni-NTA column (ThermoFisher). The Ni-NTA column was washed with 150 mL of lysis buffer with graded increases of imidazole up to 50 mM. Finally, protein that remained on the column was eluted from the Ni-NTA resin using a lysis buffer containing 300mM imidazole.

To produce a more extensively purified preparation for the dimerization assays, TelT was cloned with C-terminal 3C-StrepII-His_6_ tag (pET29b backbone, Supplementary Table 1). The expression plasmid was transformed into E*. coli* BL21 AI cells and cultured in LB with 50 μg/ml kanamycin at 37°C. At an OD600 of 1.0-1.1, expression was induced with 0.5 mM IPTG and 0.2% arabinose. Protein was expressed overnight at 16 °C. Cells were lysed by sonication in Buffer A (50 mM Tris pH 7.5, 1 M NaCl, 25 mM imidazole, 5 mM ß-mercaptoethanol, with cOmplete EDTA-free protease inhibitor cocktail (Roche), and 1 mM PMSF). The lysate was cleared by centrifugation at 72,000 g at 4°C for 30 min. The protein was purified via affinity chromatography by applying the soluble fraction to a Strep-Tactin column (StrepTrap XT; Cytiva), followed by elution with 50 mM biotin in Buffer A. Fractions were concentrated using 30 kDa molecular weight cut-off centrifugal concentrator (Vivaspin 15, Sartorius), then applied to Superdex 200 10/300 G size-exclusion chromatography column (Cytiva) equilibrated with 50 mM Tris pH 7.5, 1 M NaCl, 1 mM tris(2-carboxyethyl)phosphine (TCEP) and 5% glycerol. TnsP was expressed with cloned with N-terminal TwinStrep-SUMO-tag into expression plasmid (pXT130 backbone, supplemental Table S1), transformed into *E. coli* BL21 cells and cultured in LB media with 100 μg/ml ampicillin at 37 °C. At an OD600 of 0.6-0.8, expression was induced with 0.5 mM IPTG. Protein was expressed overnight at 16 °C. Cells were lysed by sonication in Buffer A (100 mM Tris pH 8.0, 1 M NaCl, 5 mM ß-mercaptoethanol) with cOmplete EDTA-free protease inhibitor tablet (Roche), 1 mM EDTA and 1 mM PMSF. The lysate was cleared by centrifugation at 72,000 g at 4 °C for 35 min. The protein was purified via affinity chromatography by applying the soluble fraction to a Strep-Tactin column (StrepTrap HP; Cytiva), followed by elution with 2.5 mM d-desthiobiotin in Buffer A. Part of the elution fractions were incubated with SenP2 protease in 1:100 w/w ratio overnight. Sample was further purified by Ni-affinity chromatography column (HisTrap HP, Cytiva) equilibrated with Buffer A supplemented with 25 mM imidazole. Protein was eluted with an imidazole gradient reaching 500 mM in 20 column volumes. Fractions were concentrated using 3 kDa molecular weight cut-off centrifugal concentrator (Vivaspin 15, Sartorius).

### Protelomerase activity assays

The protelomerase reaction followed the protocol of NEB TelN. Substrate DNA is incubated with protelomerase in ThermoPol reaction buffer at 30°C for 30 minutes, followed by 75°C for 5 minutes. For transposon-associated protelomerase TelT, proteinase K was added after the reaction and incubated for another 30 min at 30°C to remove tightly bound proteins from DNA. The DNA products were examined with SDS PAGE, denatured or non-denatured. All substrates used in the assay are indicated in Supplementary Table 1.

For protelomerase activity assays with short DNA substrates, DNA was mixed with TelT in 1 µM and 1.2 µM concentration, respectively, in TelT reaction buffer (20 mM Tris pH 8.0, 100 mM NaCl, 2 mM MgCl_2_, 1 mM TCEP), followed by incubation at 30 °C for 1 hour. For the control reactions, DNAs were incubated with 1 µl EcoRI in Buffer 2.1 (NEB). Reaction mixtures were treated with Proteinase K for 30 minutes at 55°C and products were separated by electrophoresis on 1xTBE 15% polyacrylamide gels, (80 minutes at 120 V). Results were visualized by ethidium-bromide staining, with UV light using EBOX VX5 gel imaging system (Vilber).

### Covalent intermediate assay

10 µM DNA substrates were combined with 1.5 µM TelT in TelT reaction buffer and incubated for 1 hour at 30 °C. The samples were concentrated ∼3-4-fold using SpeedVac; reaction products were separated by SDS-PAGE electrophoresis on 12% acrylamide SDS-PAGE gels (170 V for 1 hour) and visualized using Coomassie brilliant blue dye.

### TelT dimerization assays using EMSA

Oligonucleotides were synthesized by Integrated DNA Technologies (IDT) and purified via HPLC. The short hairpin DNA substrate TelT_short_hairpin_2_tye705 substrate was TYE™705-labeled, and the long hairpin substrate was assembled by mixing two oligonucleotides: IRDye®800-labeled TelT_long_hairpin_1_ird800 and TelT_long_hairpin_2. Hairpin formation was prompted by heating to 95°C for 5 minutes, followed by rapid cooling on ice. Sequences of all oligonucleotides are presented in Supplementary Table 1.

DNA binding and TelT oligomerization were assessed by Electrophoretic Mobility Shift Assays (EMSA). 1 µM DNA with or without 2 µM TelT was incubated in 20 mM Tris, 133 mM NaCl, 2 mM MgCl_2_, and 1 mM TCEP buffer. Reaction products were separated on 2% agarose gel electrophoresis, in 0.5x TBE (90 min, 120 V). The gel was imaged using Sapphire FL Bimolecular imager (Azure Biosystems).

### TelT dimerization assays using mass photometry

Measurements were performed with the TwoMP instrument (Refeyn) at room temperature. On a glass coverslip, silicone gaskets with 2 x 3 wells were positioned for holding the sample droplets. The instrument was calibrated using bovine serum albumin. Purified TelT was diluted to 10 µg/ml concentration in buffer containing 20 mM Tris (pH 8.0), 133 mM NaCl, 2 mM MgCl_2_, 0.5 mM TCEP. To an aliquote of the protein, 62 bp hairpin DNA substrate mimicking the LoPAST telomere end (long hairpin, see DNA substrates section) was added. The samples (2 µl) were loaded onto the glass coverslip by mixing into a buffer droplet (18 µl) and mass measurements were recorded. Data analysis was performed using the Discover MP software.

### TnsP DNA binding by EMSA

DNA oligonucleotides for EMSA were synthesized and purified via desalting by Microsynth. Oligonucleotides were annealed in Milli-Q water, or in 5 mM NaCl, 1 mM MgCl_2_ solution by heating up to 95°C for 5 minutes. Hairpin DNA substrates were quickly transferred to ice, while dsDNA substrates were cooled down gradually to promote proper annealing. Purified TnsP was incubated with 1 µM DNA substrates in 2:1 molar ratio in buffer containing 20 mM Tris (pH 8.0), 100 mM NaCl, 2 mM MgCl_2_, 1 mM TCEP for 30 minutes on ice. Samples were mixed with 5x Green GoTaq® Reaction Buffer (Promega), used as loading dye, and loaded on 1xTBE 10% polyacrylamide gels, next to Low Molecular Weight DNA Ladder (NEB). Gels were run at 120 V for 1 hour, then stained with GelRed® nucleic acid staining solution. Gels were imaged with UV light using EBOX VX5 gel imaging system (Vilber).

### Mating-out transposition assay

To assess LoPAST transposition activity, three vectors were constructed (Supplementary Table 1). The donor plasmid had a kanamycin resistance (KanR) gene flanked on both sides by the single transposon end associated with the LoPAST element in cyanobacteria (pOPO606). The TnsB, TnsC, and TniQ proteins from LoPAST were expressed as their native *tnsBCQ* operon from a Lac promoter (pOPO705). The small open reading frame encoding the protein we call TnsP from LoPAST was expressed with the Ara pBAD promoter (pOPO599). Vectors were maintained with kanamycin (pOPO606), chloramphenicol (pOPO705), and ampicillin (pOPO599), or empty vector controls. Transposition frequency was monitored in a large pool of independent transformants^16^. *E. coli* strain BW27783 with the F-*lacZ* mobilizable target plasmid and the pOPO606 plasmid encoding the *tnsBCQ* operon was freshly transformed for each experiment with pOPO705 and pOPO599, or their derivatives as indicated for each experiment. After overnight growth, hundreds of transformants were washed from the plate using LB media, pelleted, subjected to two washes with M9 minimal media, and re-suspended to an optical density (O.D.) of 0.6 in M9 minimal media. M9 minimal media had 0.2% w/v maltose, and 0.2% w/v arabinose and 0.1 mM IPTG for induction. After an 18-hour incubation period with agitation at 30°C, a 0.5 ml portion of the donor cells was pelleted, washed twice with LB, and was re-suspended in 0.5 ml LB supplemented with 0.2% w/v glucose for recuperation with agitation at 37°C for 30 minutes. To track transposition from the donor plasmid to the F plasmid target, donor cells were mixed with mid-log recipient cells (CW51) in LB supplemented with 0.2% w/v glucose at a 1:5 ratio of donor to recipient. The mixture was incubated with gentle agitation for 90 minutes at 37°C to facilitate mating. After incubation, the cultures were vortexed, placed on ice, serially diluted in LB 0.2% w/v glucose, and plated on LB supplemented with the necessary antibiotics for selecting CW51 recipient cells for transconjugants (20 μg/ml nalidixic acid, 100 μg/ml rifampicin, 50 μg/ml spectinomycin, 50 μg/ml X-gal), with or without 50 μg/ml kanamycin to either sample the entire transconjugant population or specifically select for transposition, respectively. The plates were incubated at 37°C for 24 hours before counting the colonies.

To map the insertions on the target F plasmid, colonies were washed off the plates using LB with 0.2% (w/v) glucose, washed twice with LB, and then diluted to O.D. 600 of ∼0.6 in 5 mL of LB with 0.2% (w/v) glucose. After approximately 3 hours of incubation at 37°C with agitation, the cells were pelleted, and the F plasmids containing insertions were extracted using the ZR BAC DNA Miniprep Kit. The DNA was submitted to SeqCenter for Illumina next-generation sequencing (NGS). Reads with transposon ends were processed and mapped with BBtools (<u>sourceforge.net/projects/bbmap/</u>).

A slightly modified procedure was used for testing the transposition of a type I-E CAST system from *Streptomyces* sp. NRRL S-1831. The donor plasmid had the kanamycin resistance gene flanked by the same I-E CAST transposon end sequence (pOPO1098). TnsBC from the type I-E CAST system was codon-optimized and expressed with a Lac promoter (pOPO1127). Walker B motif mutated derivatives TnsBC(E134A) (pOPO1134), TnsBC(E134Q) (pOPO1135), and TnsBC(E134K) (pOPO1137) were also tested. The only differences from the methods used in the LoPAST system is that the arabinose was not used to induce expression, only leaky expression from the vectors and transposition was monitored at 16°C. Additionally, 1 ml of the induced culture was collected for washing, recovery, and conjugation, instead of 0.5 ml.

### Conditional plasmid-based transposition assay

LoPAST transposition was monitored in *E. coli* BW27783 comparing the normal circular chromosome configuration and a linear chromosome derivative to test for any changes in the frequency or distribution of transposition events. An *E. coli* strain with a linearized chromosome was constructed based on the study of Cui, Tailin, et al^17^. To construct this strain, a DNA fragment containing the N15 phage linear end resolution site (*tos*) and a kanamycin resistance gene was first integrated into the *E. coli yneO* gene located in the replication terminus region of *E. coli* BW27783 by Red recombination. The Red recombination function was expressed in *E. coli* BW27783 from plasmid pKD46 with arabinose induction^37^. Then a DNA fragment containing *telN* and Gentamicin resistance gene is used to replace the kanamycin resistance gene. The temperature-sensitive pKD46 is cured from the resulting bacterium by culturing at 42°C, creating BW27783 *yneO*::GmR-*telN*-*tos* (PO621). Chromosome linearization was confirmed by NGS reads displaying loop-back patterns around the hairpin ends.

A curable donor plasmid (pOPO1240) is constructed for the assay. The donor plasmid backbone contains a temperature-sensitive replicon and a codon-optimized *pheS\** gene (T255S, A309G) under the J23119 promoter, which confers sensitivity to 4-chlorophenylalanine (4CP) in LB medium^38^. In addition to the *kanR* gene, a γδ resolvase gene and its resolution site from the F plasmid were incorporated into the mini-LoPAST transposon to facilitate cointegrate resolution (Supplemental table 1).

For the transposition assay, *E. coli* BW27783 and PO621 strains carrying pOPO1240 were transformed with either pOPO705 (LoPAST TnsBCQ) and pOPO599 (LoPAST TnsP), or pOPO705 and pBAD322. To monitor transposition into the chromosome the donor element with two ends was on a plasmid that had temperature sensitive DNA replication and encoded the *pheS*\* gene which confers sensitivity to 4-chlorophenylalanine (4CP) in LB medium^38^. In the assay transposon functions are expressed at the permissive temperature for plasmid replication in LB without 4CP. The plasmid with the mini element is then cured by growth at the nonpermissive temperature for plasmid replication and plated on selection plates containing 4CP which is toxic to cells with the *pheS*\* gene.

Transformants were initially selected for the assay at 30°C overnight on LB 0.2% glucose with antibiotics. Colonies were washed up from the plates with 5 ml M9 minimal medium, spun down, and washed twice with M9 medium, then resuspended into triplicates of 3 ml induction medium (M9 medium, 0.2 % Maltose, 0.2 % Arabinose, 0.1 mM IPTG, 100 μg/ml carbenicillin, 30 μg/ml chloramphenicol, 50 μg/ml kanamycin, 50 μg/ml spectinomycin). After induction at 30°C for 18 hours with agitation, 500 ul of induced culture was spun down, washed twice with LB, resuspended in 5 ml LB 0.2 % glucose, then cured of the donor plasmid by incubating at 42°C for 16 hours with agitation. The cultures were plated onto LB plates with 10 mM 4CP and kanamycin, and incubated at 42°C. A low number (∼10-20) of breakthrough colonies was detected in most transposition assay where the plasmid was not lost. These were easily detectable because they were 4CP sensitive and plasmid sequence could be identified by PCR. Colonies were counted and a subset was re-patched onto fresh plates for genomic DNA extraction and sent for total genome sequencing by NGS.

### Detecting telomeric transposons by mapping locations and orientations of DDE family transposases

DDE family transposases in *Streptomyces* are identified using two Pfam profiles, PF00665 (rve, Integrase core domain), or PF09299 (Mu-C terminal domain), utilizing the hmmsearch tool of HMMER v3.4^36^. The cutoff is set to an E-value 0.001. Only transposase genes located on assembled chromosomes were include in the analysis. Proteins smaller than 150 amino acids were excluded. The protein sequences are clustered with Cd-hit with a cutoff of 85% identity. The sequences deduplicated and then aligned using MUSCLEv5^39^. The resulting alignment is utilized to construct a protein similarity tree with FastTree^40^, employing the Whelan and Goldman (WAG) evolutionary model and the discrete gamma model (20 rate categories). The locations and orientations of transposases within the same clusters are plotted together on the tree.

### Assessing the consensus sequences of active site residues in TnsABC

Protein sequences from selected clades are gathered and deduplicated using a 95% identity cutoff with Cd-hit^41^. The sequences are aligned with MUSCLEv5^39^, and the active site residues are identified by aligning a representative with Tn7 TnsA, TnsB, or TnsC with HHpred^42^. Sequences around active site residues are extracted from multiple alignments and converted into sequence logos using WebLogo^43^.

### Detection of protospacers of type I-E CRISPR

For type I-E CRISPR associated with *cas1*-*cas2*, the DNA sequences 10 kbp upstream and downstream of the operon are collected. CRISPR spacers are detected with MinCED (https://github.com/ctSkennerton/) and clustered by 90% identity with Cd-hit^41^. For type I-E CAST, we collected 20 kbp sequences upstream of *cas3* genes for searching CRISPR. Because type I-E CAST CRISPR are often fragmented and less conserved, a different approach was taken. We identified CRISPR arrays by initially manually identifying several divergent CRISPR repeats from various type I-E CAST. These repeats were aligned using Clustal Omega^44^, and the alignment was compiled into a profile, which was then employed for CRISPR repeat detection using nhmmscan^36^. Repeats that are spaced within reasonable distances (57-65 bp) are defined as part of an array. The gathered repeats were utilized to create a profile, and this process was reiterated until consistency was achieved in repeat detection. CRISPR spacers are then clustered by 90% identity with Cd-hit. To find protospacers, spacers were BLASTn against annotated *Streptomyces* genomes. Only hits with mismatches no more than 4 nt (including non-aligned region), match protein-coding sequences, and not in CRISPR arrays were collected as protospacers.

The putative transposon ends of TnsBC telomeric transposons are identified by manually BLASTn for sequences that are consistently associated with the transposons and border different sequences.

### Data visualization

Genome visualization was conducted with the Python package pyGenomeViz 9 https://github.com/moshi4/pyGenomeViz/blob/main/CITATION.cff)^45^. Protein trees were visualized with iTOL^46^ and the Python package ETE Toolkit^47^.

## Disclosure Statement

Cornell University has filed patent applications unrelated to work in this manuscript with some of the authors as inventors.

## Acknowledgments

Work in this manuscript was funded by the NIH Grants GM129118 (JEP) and GM152260 (JEP), the Canton of Geneva (OB) and European Research Council ERC-2019-COG project mARs, grant agreement number 866250. (OB).

---

**Figure S1.**
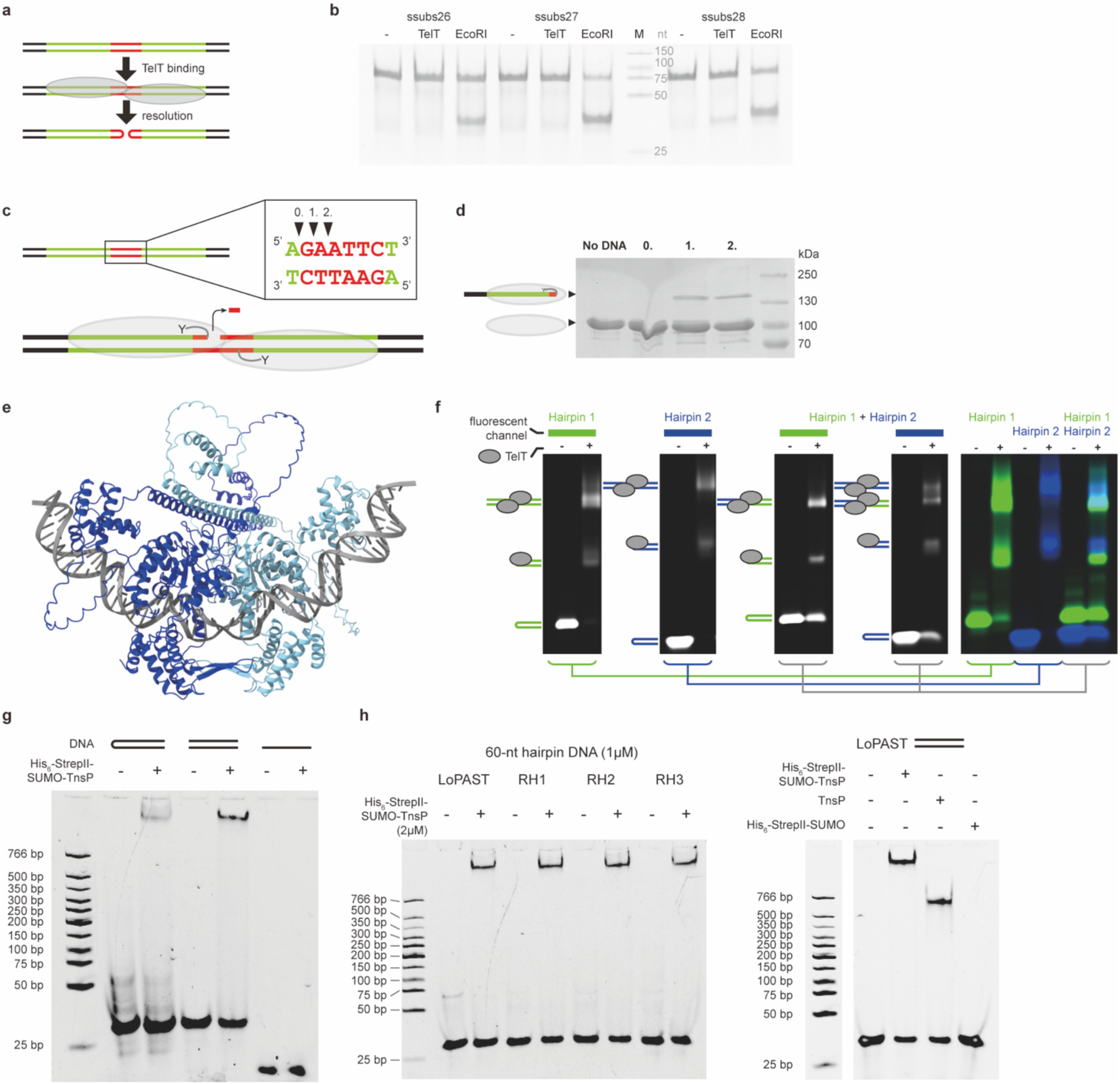
DNA requirements for TelT activity, TelT follows the canonical protelomerase biochemical pathway. (a) Scheme of telomer resolution. The short DNA substrates (ssubs) used in the assay are color-coded: green – native telomere sequences for TelT binding, red - putative cleavage region, black – GC-rich ends to facilitate DNA annealing. (b) Telomer resolution assay shows detectable resolution of a substrate with 28 bp long native telomere sequence on native PAGE. EcoRI was used as the positive control. (c) Scheme of the DNA cleavage assay with ‘suicide’ substrates. The substrate sequences are identical to ssubs-28, with one strand nicked at positions 0., 1., or 2. (black triangles). If the nick is placed 3’ to the cleavage position, TelT activity liberates one or two nucleotides, which diffuse away trapping the covalent phosphotyrosyl protein-DNA intermediate, which can be detected by SDS-PAGE. (d) Cleavage assay results show covalent TelT-DNA intermediate with nicks at positions 1. and 2., indicating that TelT cuts DNA at position 0. (e) The AlphaFold3 web server was used to predict the supramolecular assembly of TelT homodimer (shades of blue) binding 28 bp DNA repeats^1^. Besides the core protelomerase domains a C-terminal extension is predicted to bind the outer DNA segments. (f) The addition of TelT to fluorescently labelled telomere end-representing hairpin DNA substrates results in two shifted bands on agarose gel electrophoresis. Schemes indicate the putative complex stoichiometries. When TelT is added to a 1:1 mixture of two different-length substrates with different labels, one of the shifted bands is marked with both fluorophores, indicating a complex with two different DNA substrates bridged by a TelT dimer. Complexes with two “Hairpin 1” DNAs are expected to migrate similar to other complexes and cannot be separated. Single fluorescent channels are shown separately (left) and overlaid (right). (g) Electrophoretic mobility shift assay (EMSA) of DNA fragments, with sequence identical to the LoPAST telomere end, in the form of hairpin, blunt end dsDNA, and ssDNA. Addition of TnsP results in complex formation with hairpin and dsDNA. (h) EMSA of 60-nt hairpin DNA substrates (1µM), corresponding to putative LoPAST telomere ends (LoPAST), or random sequences, with GC content of 13% (RH1), 20% (RH2) and 60% (RH3). TnsP binds DNA regardless of its GC content (left), and the purification tag does not impact TnsP binding (right).

**Figure S2.**
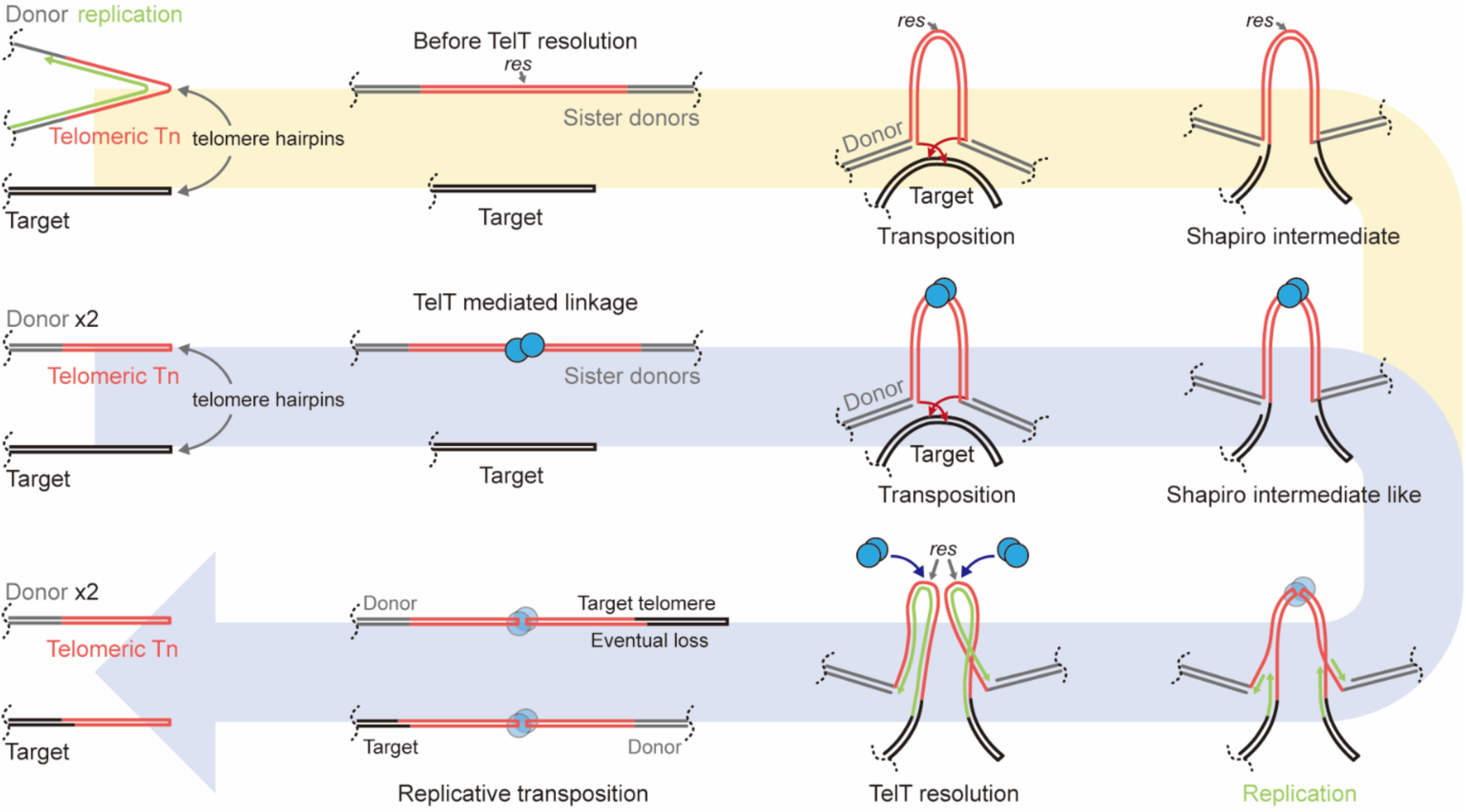
The hypothetical transposition mechanisms of PASTs. Blue path: With the help of TelT dimerization, two PASTs can be brought together, each of them provides one transposon end for transposition. Following strand-transfer, replication, and resolution, the donors are restored, and a PAST is copied and replaced the target’s telomere. Yellow path: Alternatively, DNA replication over the telomere hairpin creates an inversely repeated duplex PAST with two transposon ends before TelT resolution. Transposition of the temporary two-end PAST followed by resolution and replication results in the same replicative transposition as above.

**Figure S3.**
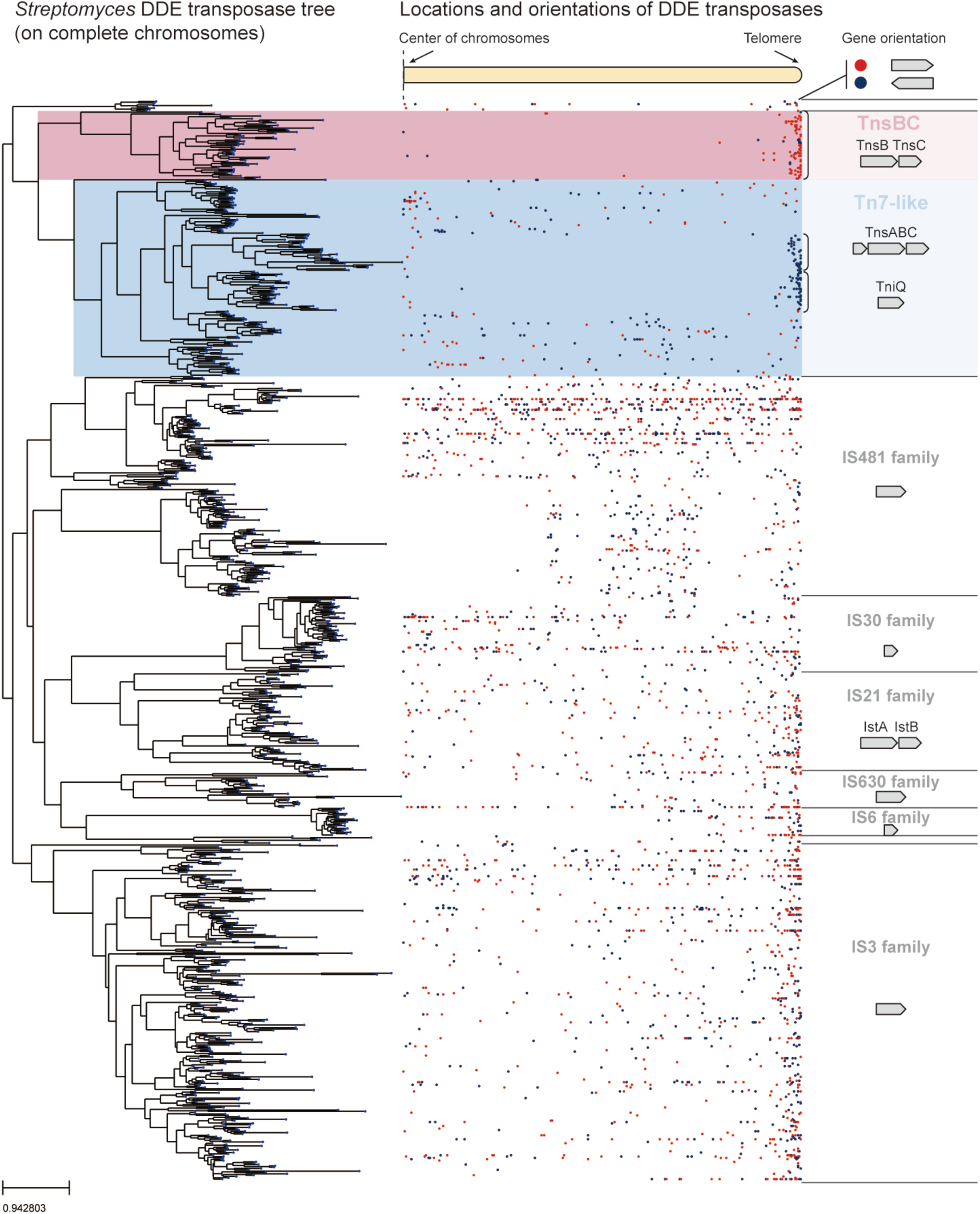
Mapping the spatial distribution and orientations of DDE family transposases along linear chromosomes in *Streptomyces*. DDE family transposases are enriched near *Streptomyces* telomeres, but the distribution varies according to transposase families. Putative telomeric transposons are located almost exclusively near telomeres and have consistent orientations. DDE family transposases were grouped using a similarity tree. The tree is made with DDE family transposases deduplicated with 85% identity as cutoff; the locations and orientations of transposases with higher than 85% identity are plotted together.

**Figure S4.**
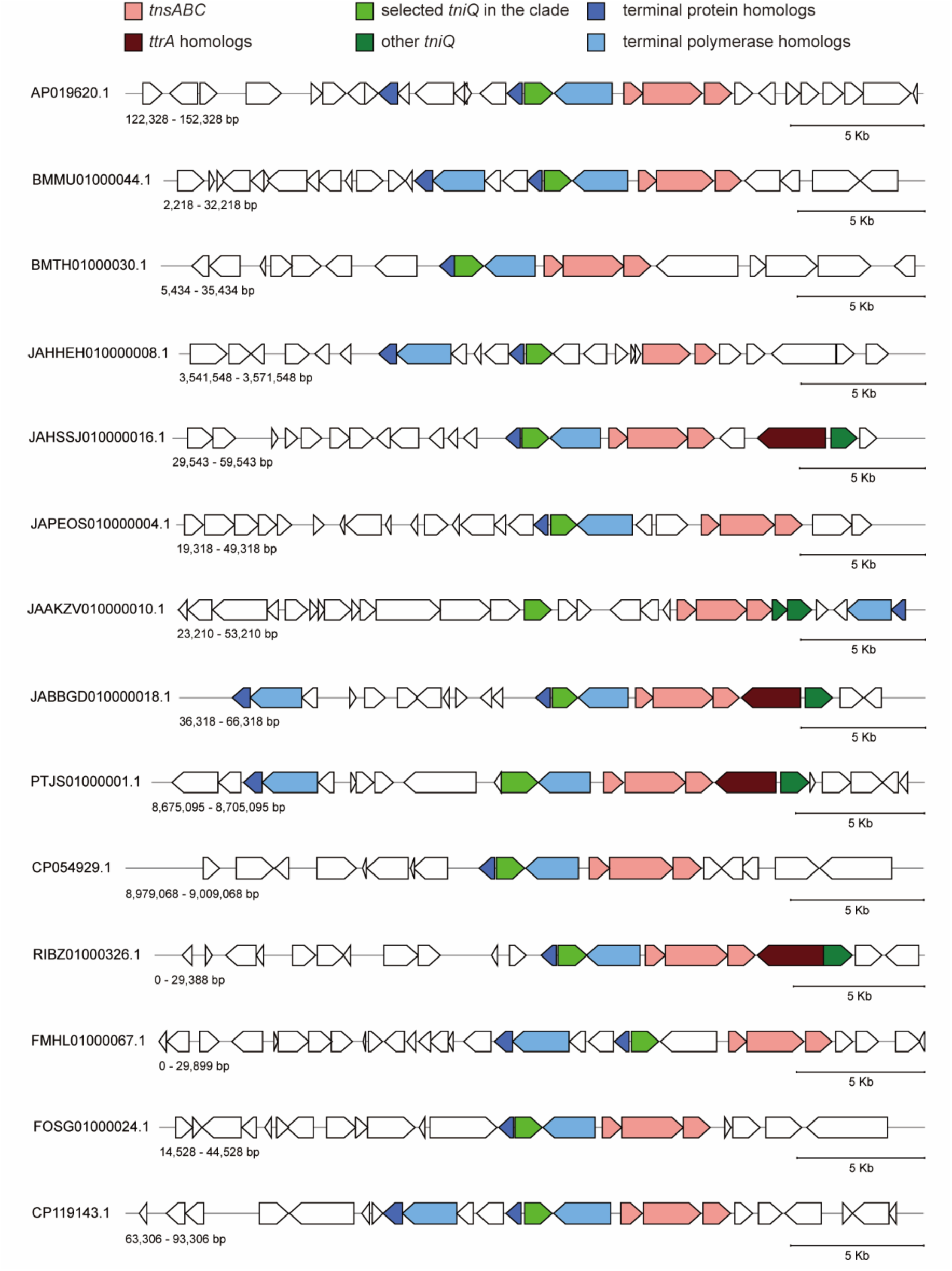
Additional examples of the genetic neighborhood of clade 1A Tin. All examples shown here have <70 % amino acid identity. The contig designation and base pair (bp) range is indicated.

**Figure S5.**
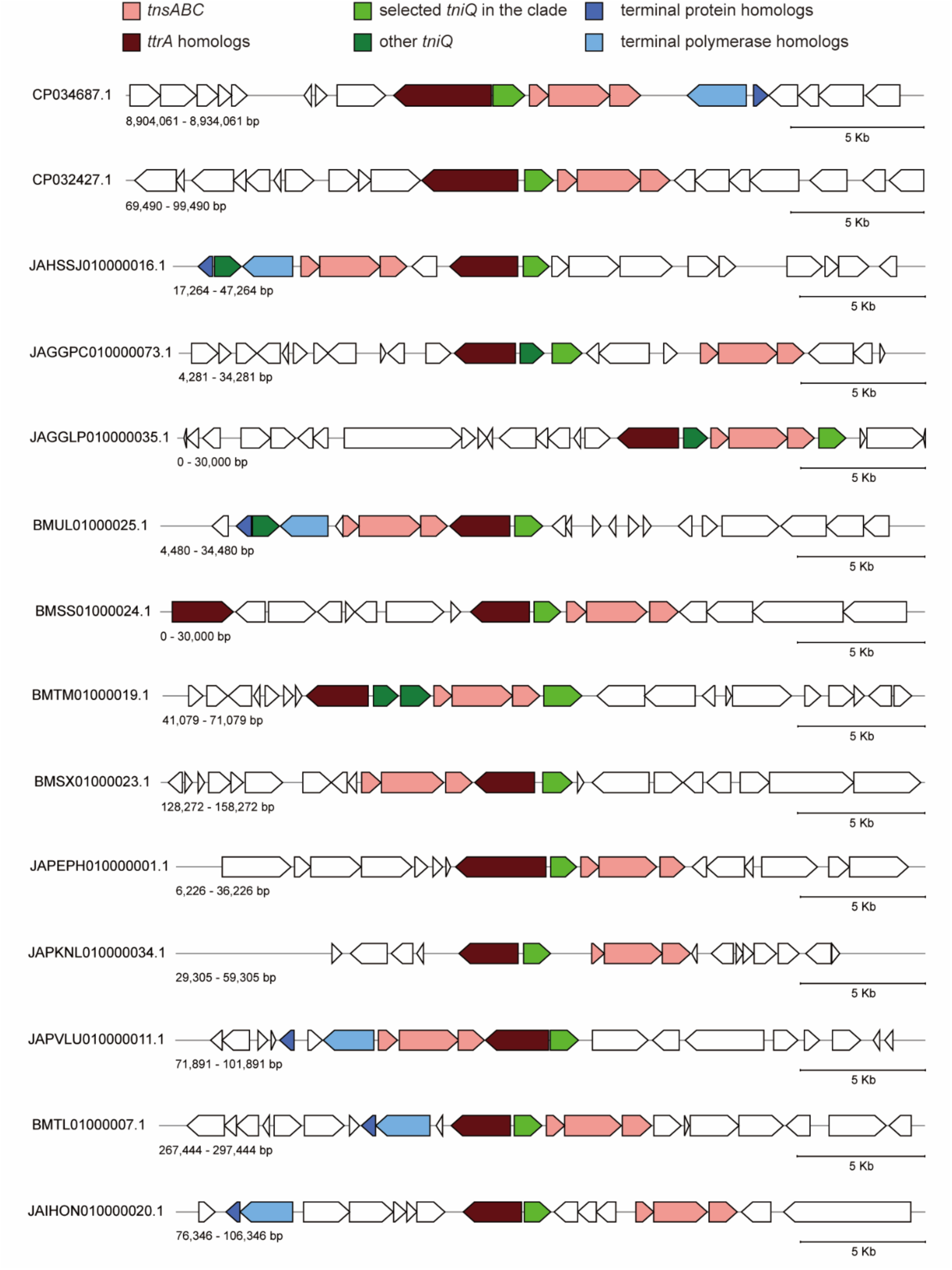
Additional examples of the genetic neighborhood of clade 1B TniQ. All examples shown here have <70 % amino acid identity. The contig designation and base pair (bp) range is indicated.

**Figure S6.**
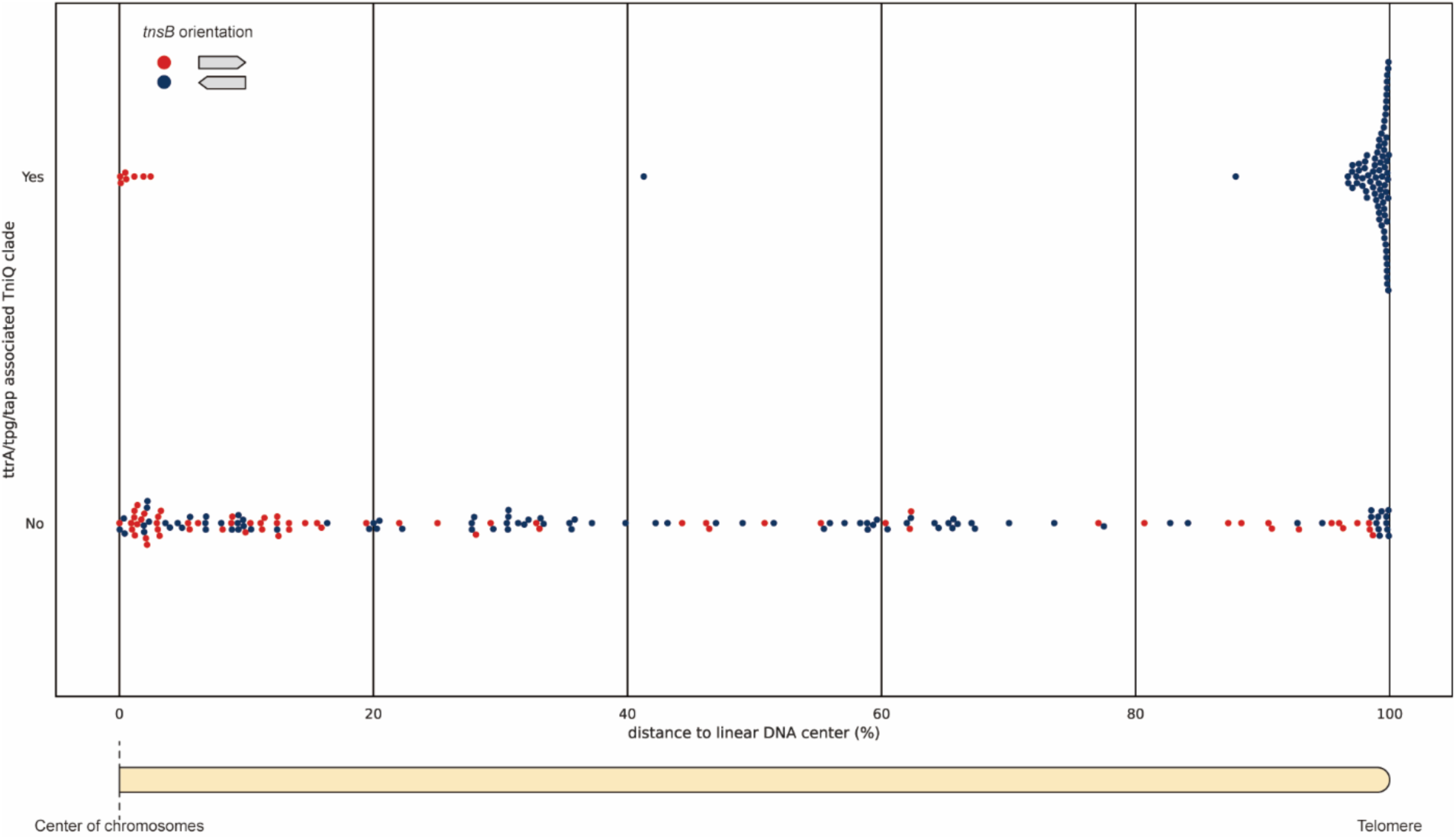
The *tnsB* genes from elements associated with *ttrA*/*tpg*/*tap* (Figure 3b colored branches) are found near telomeres in *Streptomyces* genomes. The location and orientation of the *tnsB* gene associated with TniQ branches and their association (yes) or lack of association (no) with *ttrA*/*tpg*/*tap* genes.

**Figure S7.**
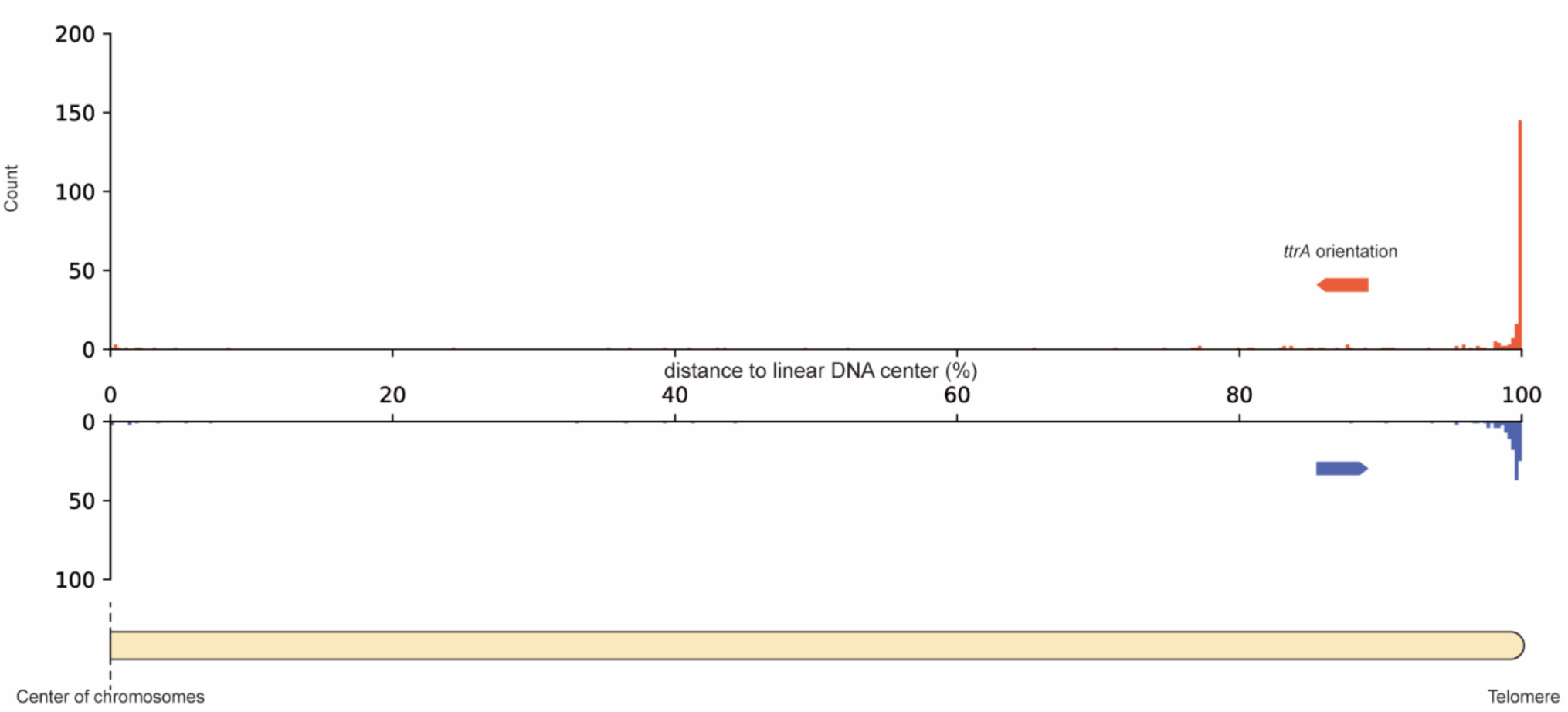
As previously observed, *ttrA* genes are primarily located at the telomeres in *Streptomyces* chromosomes. The distribution of 417 *ttrA* homolog genes on complete *Streptomyces* chromosomes is plotted according to their relative positions to the center of linear chromosomes. This aligns with previous observations that they are primarily located in the subtelomeric region.

**Figure S8.**
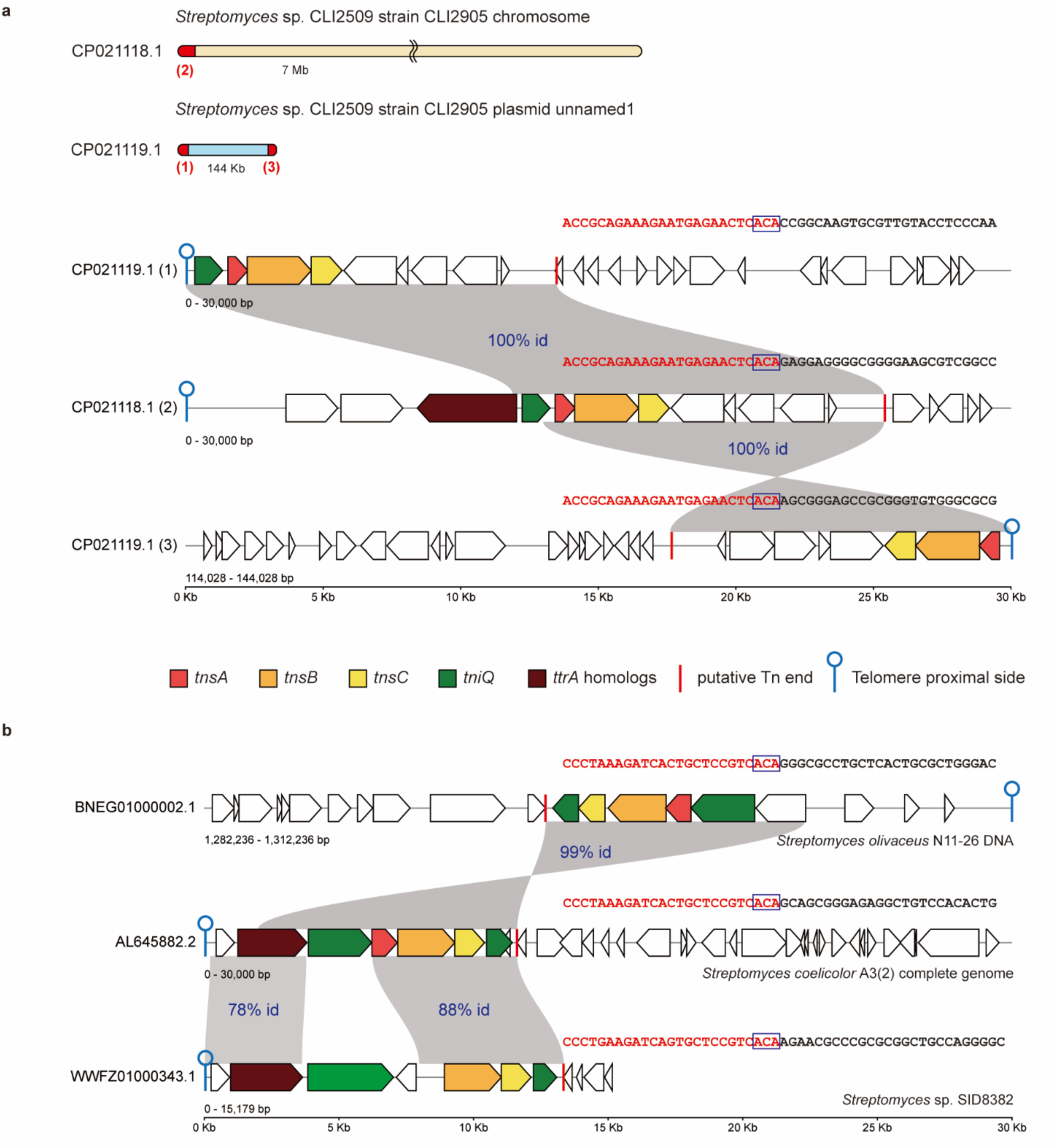
Evidence of Tn7-like telomeric transposons mobilizing across telomeres. (a) Three of the four telomeres of *Streptomyces* strain CLI2509 are populated by a single Tn7-like telomeric transposon (red). Each copy is identical, suggesting recent mobilization. The boundary of these transposons is defined by 5’-TGT/ACA-3’ sequence, typical of Tn7-like transposons. (b) Different *Streptomyces* species and strains can harbor near identical Tn7-like telomeric transposons in different locations, suggesting that the transposons can move between different hosts. The transposon boundary is also defined by 5’-TGT/ACA-3’.

**Figure S9.**
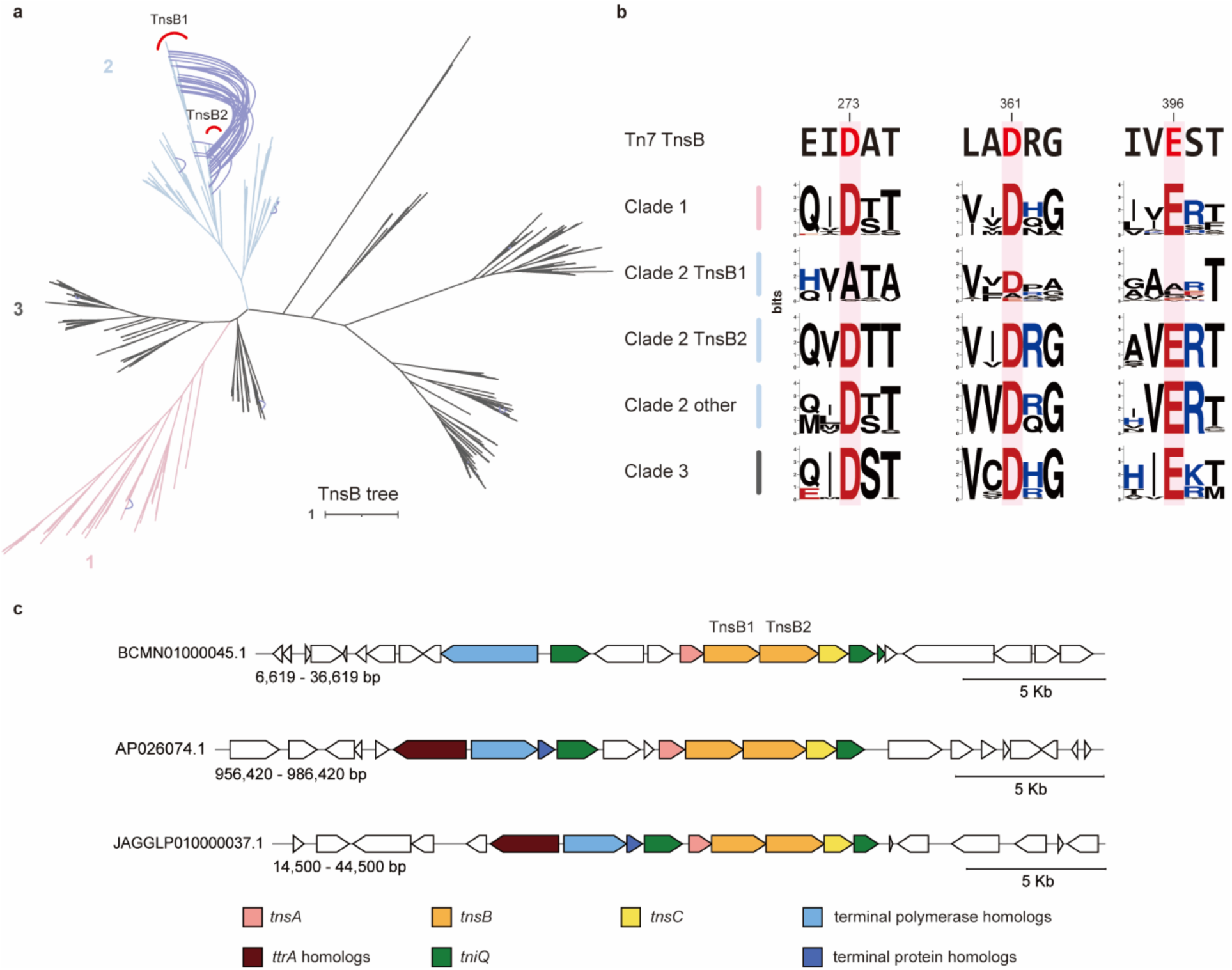
A clade of Tn7-like telomeric transposons from *Streptomyces* encodes two TnsB proteins, one predicted to be inactive. (a) Similarity tree of TnsB proteins from *Streptomyces* Tn7-like transposons. The two TnsB clades of Tn7-like telomeric transposons are colored. TnsB encoded by tandem *tnsB* genes are connected with blue lines (labeled as TnsB1 and TnsB2). (b) The amino sequence consensus near the active site residues of TnsB proteins of different transposon clades. The TnsB1 of clade 2 lost conserved active site residues needed for transposition. The active site sequences of prototypical Tn7 TnsB are indicated. (c) Genetic structure for three examples of clade 2 Tn7-like telomeric transposons with duplicated TnsB.

**Figure S10.**
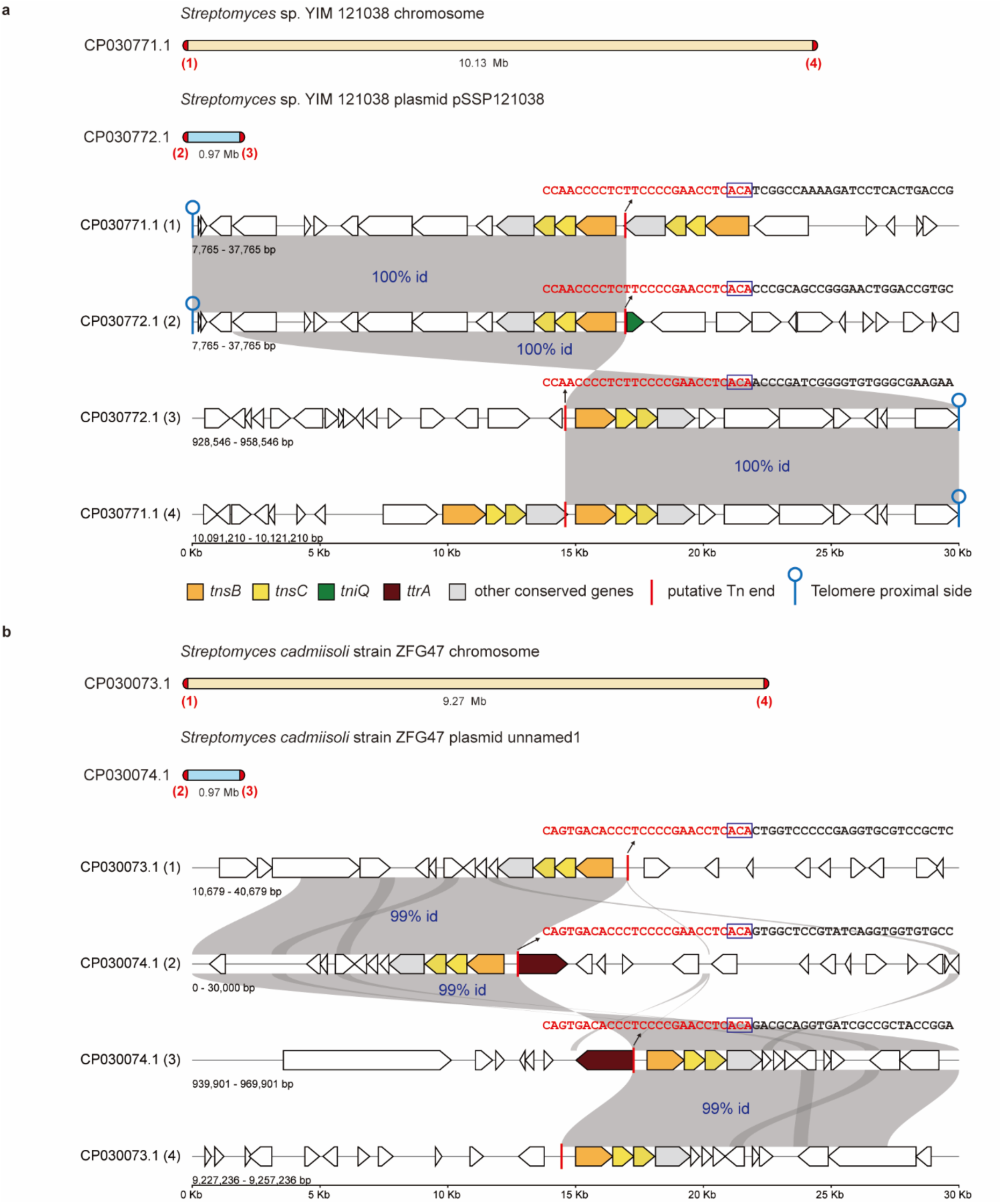
Evidence of TnsBC telomeric transposons mobilizing across telomeres. (a) A single TnsBC telomeric transposon populates all 4 telomeres of *Streptomyces* sp. YIM121038 chromosome and plasmid. Each copy is identical, suggesting recent mobilization. The boundary of these transposons is defined by 5’-TGT/ACA-3’, typical of TnsBC transposons. The chromosomal copy is inserted after another TnsBC telomeric transposon, and one of the plasmid-born copies is inserted into a *tniQ* gene of a Tn7-like telomeric transposon. (b) Similarly, in *Streptomyces cadmiisoli* strain ZFG47, a TnsBC telomeric transposon occupies all four telomeres. Two copies of the transposon disrupted *ttrA* homologs on the plasmid.

**Figure S11.**
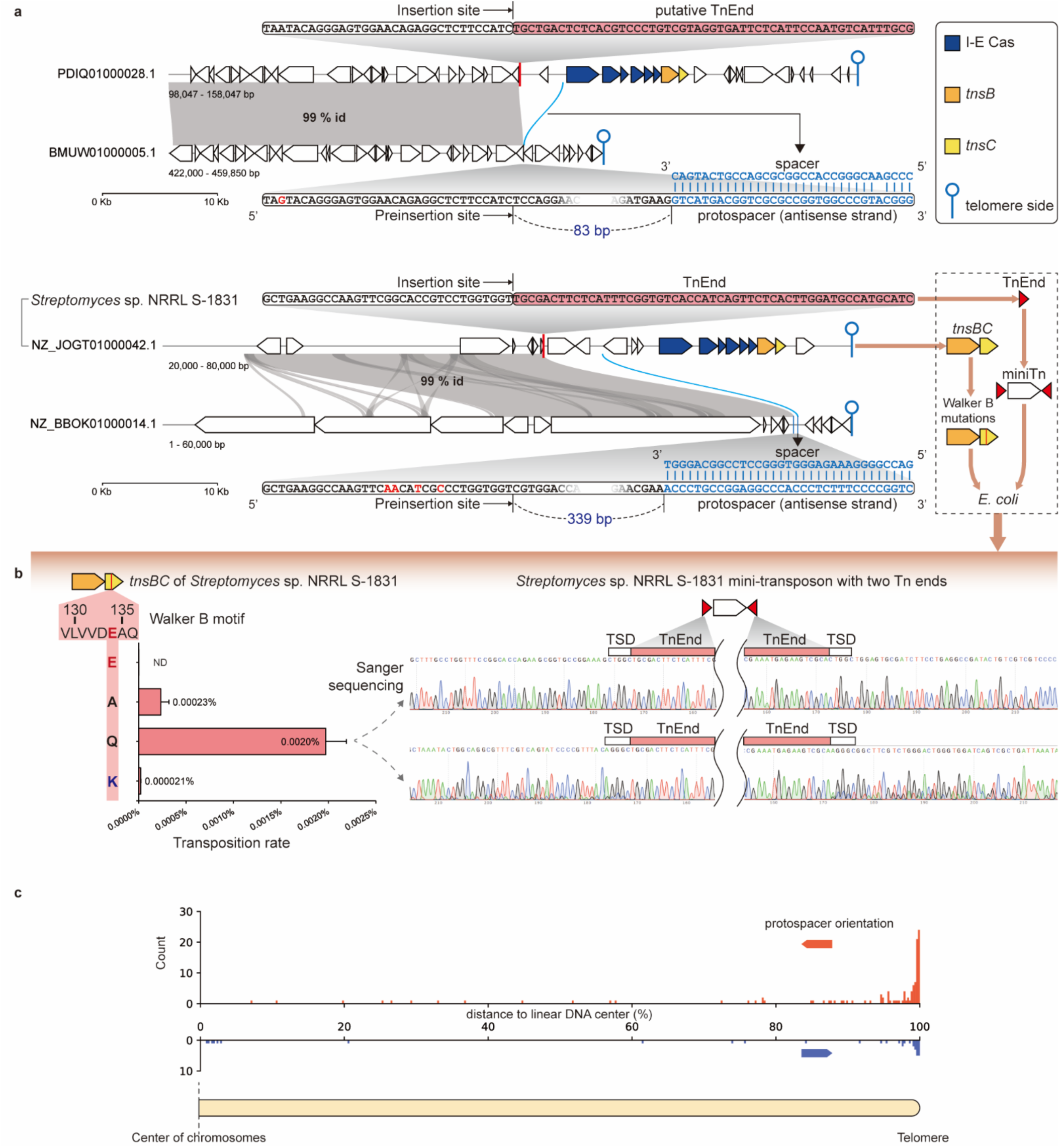
Transposition of type I-E CAST in *Streptomyces*. (a) Two examples of I-E CAST with their putative pre-insertion target DNA. Alignments are colored grey, and protospacer-spacer alignments are colored blue. The distances between protospacers and insertion positions are different in these two examples. TnEnd: transposon end. (b) Transposition assay of type I-E CAST of *Streptomyces* sp. NRRL S-1831 in *E. coli* proves that the I-E CAST is indeed a transposon, and likely requires two ends for transposition. The mini-transposon donor is made by flanking a kanamycin-resistant gene with the identified transposon end. The wildtype TnsBC shows no detectable activity in *E. coli*, likely due to the lack of natural DNA substrates, host factors, and assisting type I-E cascade. Mutations of the TnsC Walker B motif result in untargeted transposition, the mutated amino acids are illustrated on the left panel. Sanger sequencing of the insertion sites revealed the presence of 5-bp target site duplications (TSDs) flanking the two transposon ends, a characteristic signature shared with TnsBC/Tn7-like transpositions. (c) Protospacer location and orientation found with I-E CAST on complete linear chromosomes. Protospacers orientated away from telomeres are more common than protospacers oriented toward telomeres.

**Figure S12.**
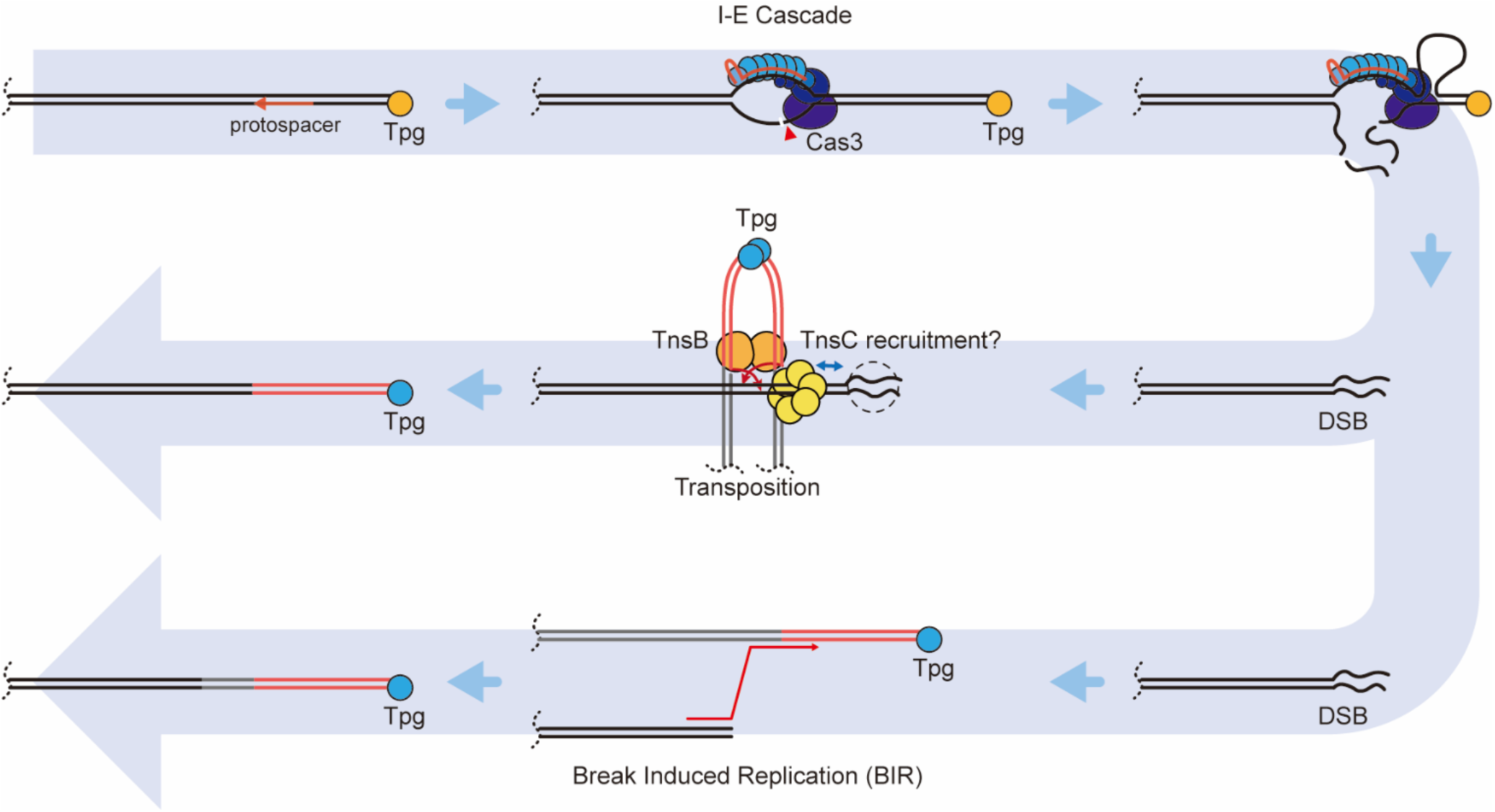
Transposition target site selection by type I-E CAST in *Streptomyces*. A proposed targeting mechanism for type I-E CAST transposition. The I-E Cascade recognizes the protospacer and activates Cas3 to nick the non-target strand, then reeling and degrading of the chromosome arm. The double-strand break (DSB) may somehow be recognized by TnsC leading to the observed transposition bias to telomeres. Alternatively, this process may promote the copying of the chromosome arm where the I-E CAST resides to the double-strand break (DSB) through break-induced repair, a mechanism known to homogenize *Streptomyces* chromosome arms.

**Figure S13.**
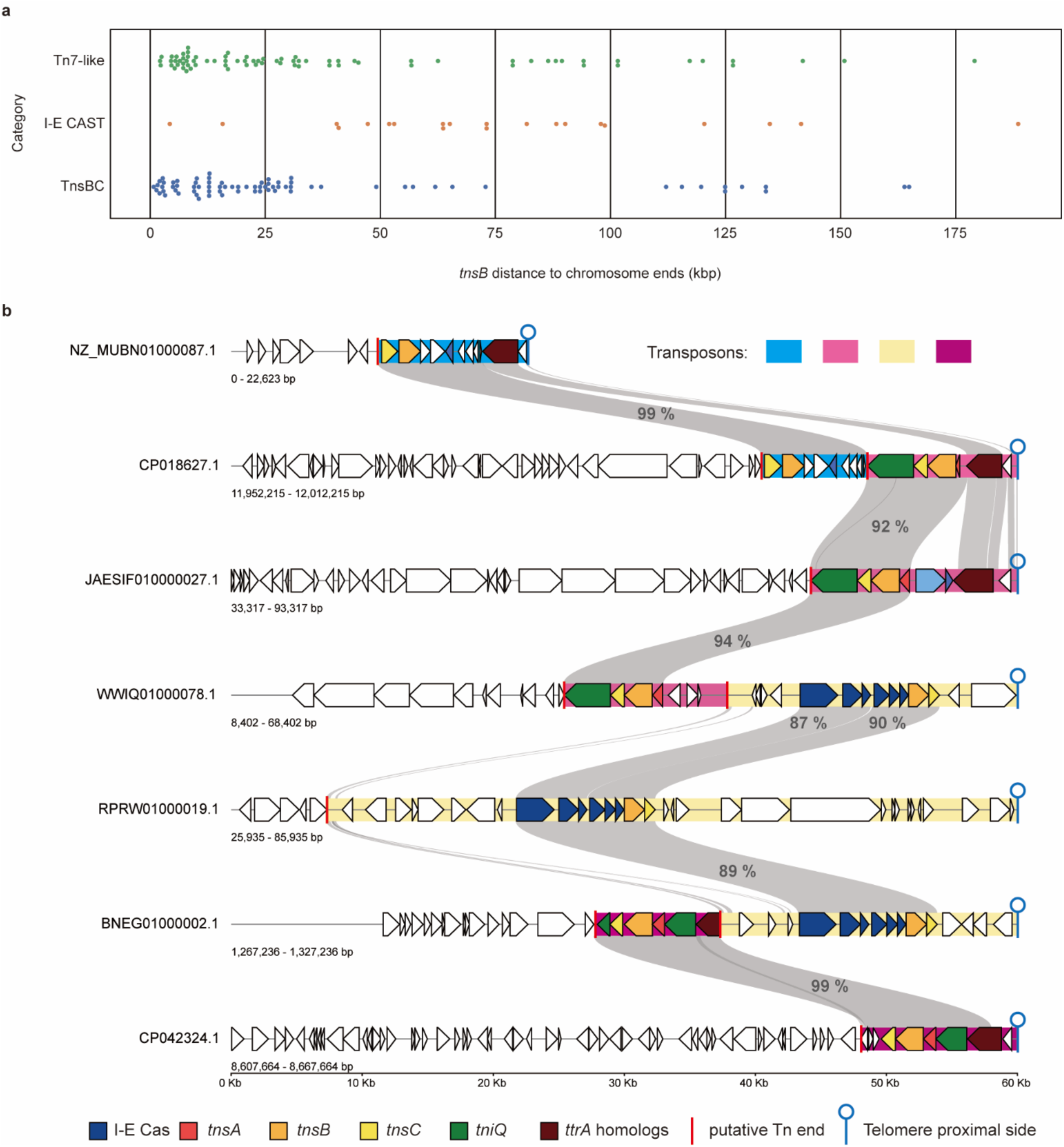
Estimating cargo size variation across different categories of transposons discussed in this study. (a) The cargo size of telomeric transposons estimated by the distance between the *tnsB* gene and the chromosome end in assembled genomes. The few examples estimated to be longer than 200 kbp are not shown. (b) Telomeric transposons contribute to telomere turnover and are predicated to compete for the same niche. Four telomeric transposons residing on seven contigs are shown. Homologous segments are connected with grey areas, nucleotide identities are labeled. Different telomeric transposons can form tandem insertions, sometimes leading to the disruption of existing elements.

**Figure S14.**
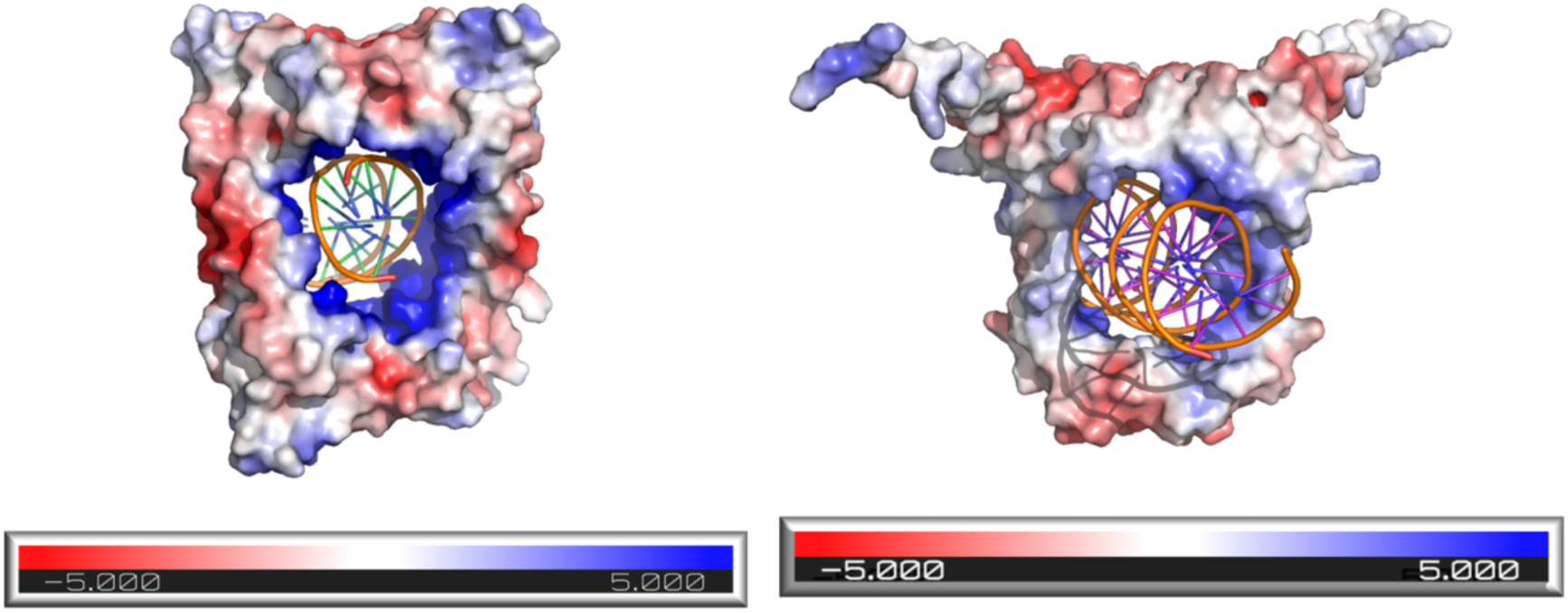
Structural models of GapR and TnsP with DNA. The homo-tetrameric GapR-DNA complex structure was retrieved from the PDB (PDB ID: 6CG8). TnsP - DNA complex was modelled using the AlphaFold3 web server^1^. TnsP is predicted to assemble a homodimer in complex with 23 bp DNA sequence (DNA sequence used to resemble a LoPAST telomere end segment). Protein surfaces are colored according to electrostatic charges (red, negative; blue, positive) using PyMOL APBS Plugin (Schrödinger, L., & DeLano, W. (2020), http://www.pymol.org/pymol)^2^. In the ring shaped TnsP model, positive surface charges surround the DNA and widen the minor groove. This configuration is consistent with the idea that the complex may preferentially interact with substates with superhelicity that are different from the bulk of the chromosome, possibly allowing the complex to recognize subtelomeric regions in linear chromosomes.

**Figure S15.**
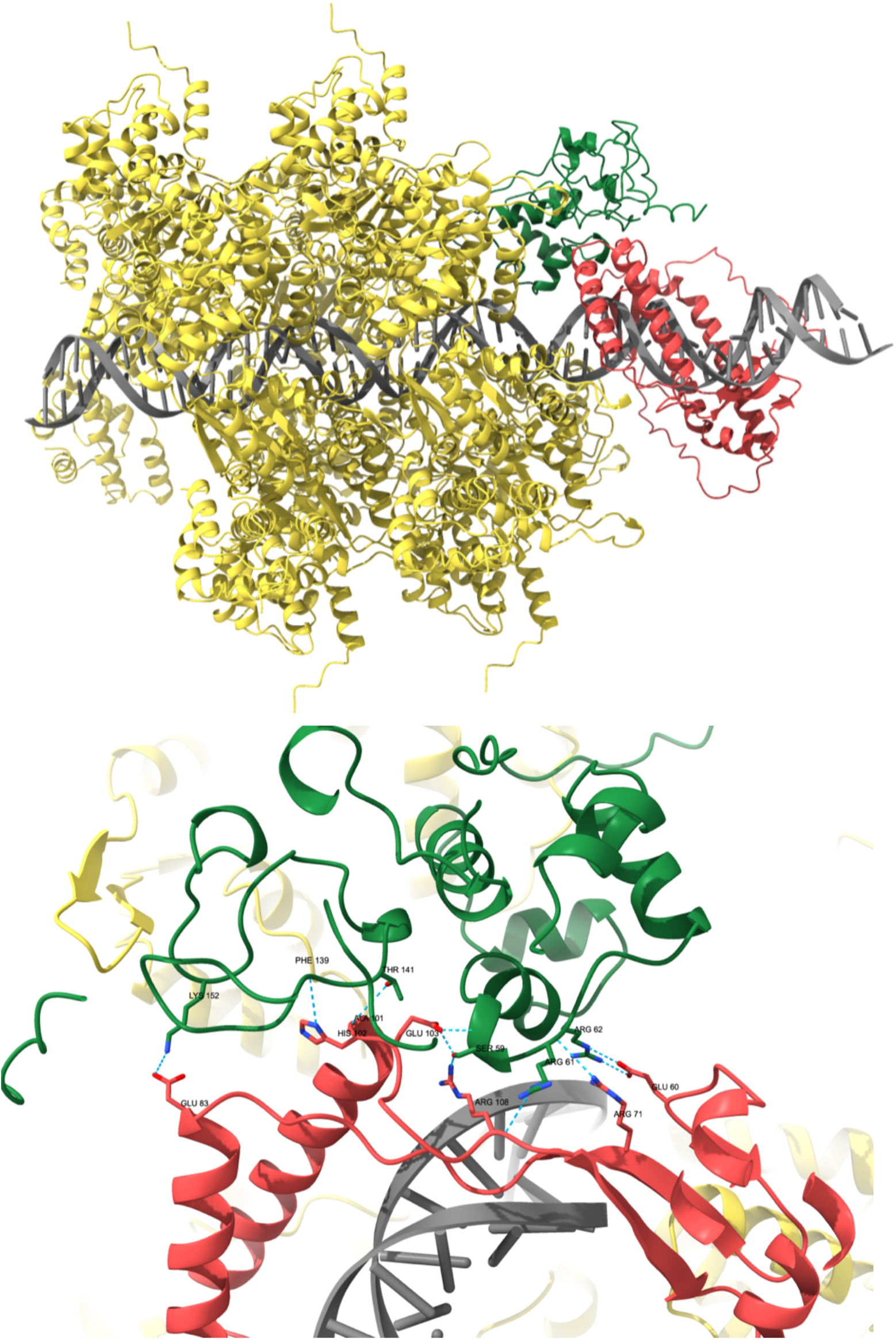
Alphafold model of TnsP in complex with other transposon proteins and DNA. The AlphaFold3 web server was used to predict the protein structures and their supramolecular assembly^1^. Images were prepared using Chimera-X^3^. The complex contains a TnsP dimer (red), a single TniQ (as in the ShCAST transpososome, 8EA3) (green), and 12 TnsC monomers (yellow). AlphaFold places the TnsP dimer on DNA, connected by TniQ to the TnsC filament. The detailed view shows the interface between TnsP (red) and TniQ (green) with H-bonds marked with the blue dotted lines. LoTniQ has a 10 residue C-terminal tail, that is not present in the ShTniQ of the CAST element and could not be predicted in the structure but could contribute to the protein-protein interactions. A 62 bp DNA sequence matching LoPAST telomere end was used in the prediction.

